# Low intensity pulsed ultrasound activates excitatory synaptic networks in cultured hippocampal neurons

**DOI:** 10.1101/2024.09.23.614451

**Authors:** Fenfang Li, Jia Wei Lin, Hao Jiang, Yu Yong, George J. Augustine

**Affiliations:** Institute of Biomedical Engineering, Shenzhen Bay Laboratory, P.R. China, 518107; Program in Neuroscience & Mental Health, Lee Kong Chian School of Medicine, Nanyang Technological University Singapore 308232, Singapore; Temasek Life Sciences Laboratory, Singapore 117604, Singapore

**Keywords:** Ultrasound neuromodulation, synaptic transmission, electrophysiology, calcium imaging, neuronal networks

## Abstract

Ultrasound can non-invasively penetrate deep into brain for neuromodulation and has demonstrated good potential for clinical application. Excitation or inhibition of neurons by ultrasound has been reported, but the underlying mechanisms are largely unknown. So far most in vitro studies have focused on the activation of individual neurons by ultrasound with calcium imaging. As the focal region of ultrasound is typically millimeter or submillimeter size, it is important to investigate yet so far unclear how the mechanical effects of ultrasound would influence on the synaptic circuit activity of neurons.

**Methods:** Low-intensity pulsed ultrasound (LIPUS) (25 MHz, 5% duty cycle, 5 Hz pulse repetition frequency, 0.4 – 1.6 W/cm^2^) was used to stimulate cultured hippocampal neurons. Action potentials and excitatory postsynaptic currents were recorded in individual cells with the whole-cell patch-clamp technique. We also simultaneously imaged intracellular calcium, along with neuronal electrical signals, to resolve neuronal network dynamics during LIPUS.

**Results:** Excitatory postsynaptic currents (EPSCs) were evoked by LIPUS in high-density neuronal cultures. Both the frequency and amplitude of EPSCs increased, indicating enhanced glutamatergic synaptic transmission. The probability of evoking responses, as well as the total charge of EPSCs evoked by ultrasound, increased with ultrasound intensity. Mechanistic analysis reveals that extracellular calcium influx, action potential (AP) firing and synaptic transmission are necessary for the responses to ultrasound in the high-density culture. In contrast, EPSCs were not enhanced in cultures with low densities of neurons. Simultaneous calcium imaging of neuronal network activity indicated that recurrent excitatory network activity is recruited during ultrasound stimulation in high-density cultures.

**Conclusion:** Ultrasound can activate recurrent neuronal network activity, caused by excitatory synaptic transmission, over tens to hundreds of seconds. Our study provides insights into the mechanisms involved in the response of the brain to ultrasound and illuminates the potential to use ultrasound to regulate synaptic function in neurological disorders that involve synaptic dysfunction, such as Parkinson’s disease and Alzheimer’s disease.

**Graphical Abstract:** 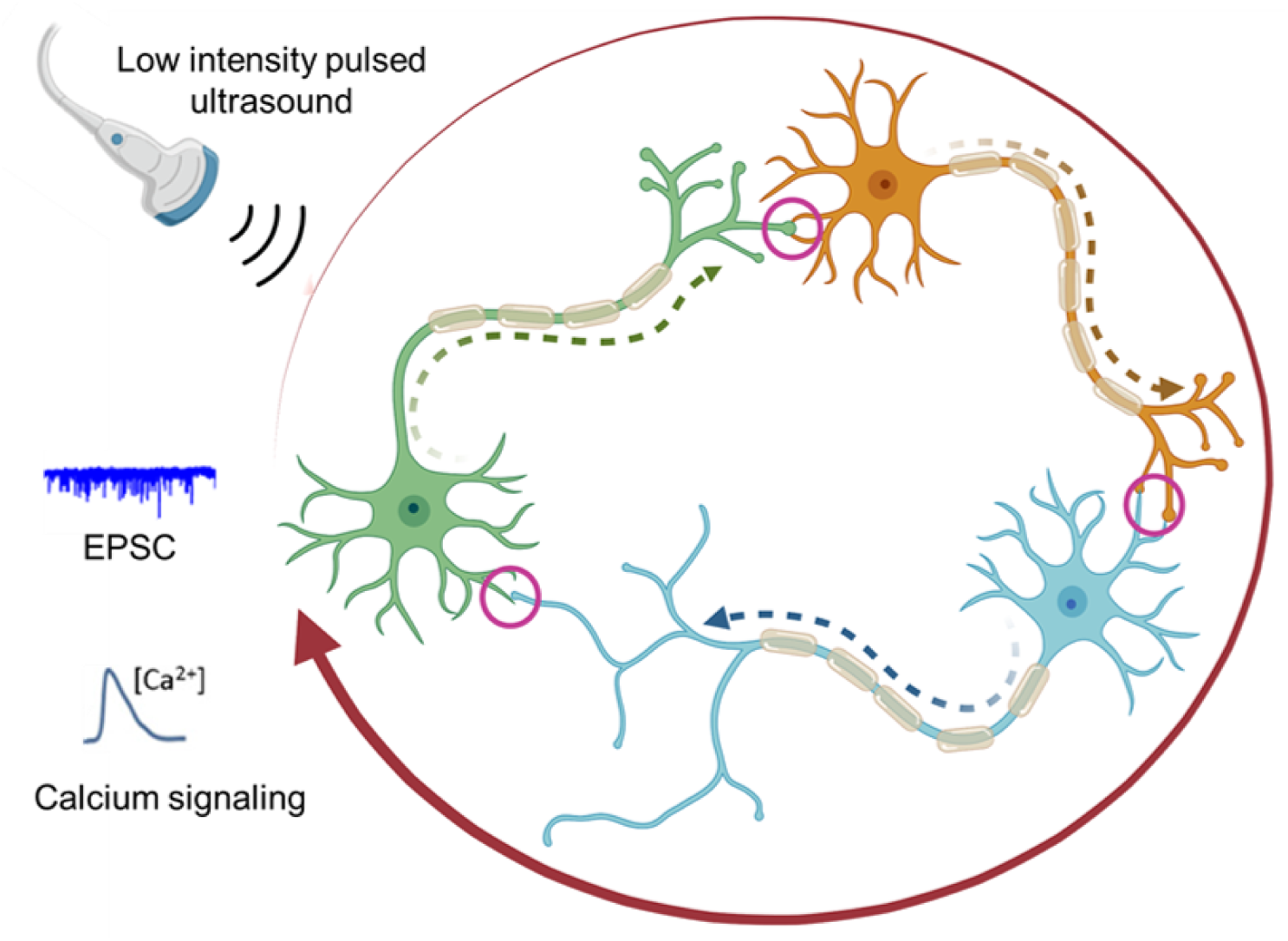

## Introduction

The development of non-invasive neuromodulation techniques has provided both the means to investigate intrinsic brain functions, as well as therapeutic strategies to modulate abnormal brain activity in neurological and psychiatric disorders. Two well-established non-invasive approaches are transcranial electric stimulation and transcranial magnetic stimulation. However, these both suffer from limitations in their spatial resolution and penetration depth [1]. In contrast, focused ultrasound (FUS) can non-invasively target deep brain regions with millimeter or better spatial precision without affecting cells along the propagation path [2]. Without genetic or chemical manipulations, and without side effects, transcranial low-intensity FUS has successfully elicited neural activity in different brain regions and behavioral responses in small animals [3–7], large animals [8–10], non-human primates [11–14] and human subjects [15–17]. These advantages make FUS promising both for basic neuroscience research and for treatment of brain disorders.

Though there is widespread interest in FUS technology, the mechanisms underlying the ability of FUS to stimulate brain activity in vivo are not well understood. FUS may work by activating mechanosensitive ion channels [18–20], transiently creating pores in the plasma membrane [2, 20, 21], or by actuating intramembrane cavitation that works via a bio-piezoelectric mechanism [22–24]. Ultrasound also affects synaptic transmission [25–27]. In mouse hippocampal slices, low-intensity ultrasound triggers exocytosis mediated by SNARE proteins and thereby enhances synaptic transmission[25]. Low-intensity FUS stimulation of the rat thalamus has been found to increase extracellular concentrations of the neurotransmitters serotonin (5-HT) and dopamine, consistent with FUS stimulating synaptic activity [26]. In a recent study, low intensity ultrasound stimulation was shown to enhance dopamine release in the striatum of a Parkinsons disease mouse model and restore the locomotor activity of these mice [27]. However, it is challenging to determine whether such mechanisms are involved in FUS stimulation of brain activity in an in vivo context; this challenge is increased by recent findings of auditory confounds during FUS use in rodents [28–30].

In this study, we used cultured hippocampal neurons to examine the actions of FUS on synaptic transmission and synaptic network activity. We found that a brief bout of FUS triggers a prolonged barrage of excitatory synaptic activity. The magnitude and duration of such responses increased with higher FUS power. Because these excitatory responses could trigger action potentials in the hippocampal neurons, we hypothesized that recurrent excitation of the hippocampal neurons was responsible for the prolonged response to FUS. Consistent with this hypothesis, calcium imaging experiments demonstrated that FUS produced a sequential activation of many neurons, with a time course comparable to that of the prolonged barrage excitatory synaptic transmission produced by FUS. Our results indicate that ultrasound can produce sustained, recurrent excitation of neural networks and such actions may underlie the ability of FUS to stimulate the brain in vivo. These results help to bridge our understanding of how FUS excitation of individual neurons can lead to neuronal network activation in the in vivo context.

## Materials and Methods

### Hippocampal neuron cultures

Newborn pups (postnatal day 0-1) from wild-type mice were used to prepare hippocampal neurons. The procedures used to maintain and use these mice were approved by the Institutional Animal Care and Use Committee of Nanyang Technological University. High-density and low-density cultures of hippocampal neurons were prepared as described in Ref. [31] and used 10-18 days later to allow the neurons to mature. The seeding concentration into 24 wells plates was around 80,000 cells/mL for high-density cultures and 20,000 cells/mL for low-density cultures. Microisland cultures of autaptic neurons were prepared with the addition of glia feeder cells to support neuronal survival [32]. After 14-18 days, these autaptic neurons were used for electrophysiological recording.

### Ultrasound setup

Ultrasound waveforms were designed by a function generator (Rigol DG972) and gated by the TTL pulse from a digitizer (Digidata 1440A) via pClamp software. The waveforms were amplified by a 75-W RF power amplifier (E&I, model A075). A 25 MHz transducer (V324-N-SU-F0.5IN, Olympus) was used to stimulate neuron cultures. The transducer output was characterized by a needle hydrophone (ONDA HNA-0400) measurements in 0.05-mm increments in y and z directions, and 0.25 mm increments in x direction in a large tank filled with water. After finding the focal point, the output pressure was measured there at different input voltage (Fig. S1).

### Particle Image Velocimetry

Polystyrene beads (2 μm) suspension in 1× DPBS (2.6% w/v) were used as tracers to map the flow field produced by the 25MHz ultrasound transducer in our experimental setup. The movement of tracer beads in the medium were recorded using a high-speed camera (Nova S12, Photron) at 12,800 frames per second with a 78 μs exposure time and were analyzed offline using the Particle Image Velocimetry (PIV) lab in MATLAB. Pre-processing was performed to locally enhance contrast using Contrast Limited Adaptive Histogram Equalization (CLAHE) with 64 pixels window size. Fast Fourier Transform (FFT) window deformation method was employed for the PIV setting with 3 passes and interrogation areas 80 × 80 × 40 × 40 and 20 × 20 pixels with 50% overlap. The correlation robustness was set as ‘standard’.

### COMSOL Multiphysics simulation

The numerical simulation was performed in COMSOL Multiphysics 6.0 trial version. The geometry was drawn according to the experimental setup and the focal length of the ultrasound transducer, where the sound wave incidents at a 45 degree angle, relative to the glass bottom of the recording chamber, and was reflected by the bottom. The computational fluid dynamic (CFD) module, acoustic module and the acoustic-fluid coupling module were selected for modeling. The 1^st^ order acoustic fields (sound pressure and acoustic radiation force) were solved using the Thermos-viscous Acoustics interface in the Acoustics Module. The streaming flow was solved using the Laminar Flow physics interface of the CFD Module and the Acoustic-Fluid Coupling Module by adding the appropriate time-averaged, first-order sources: a mass source and a volume force. The results were obtained with transient analysis and the time step was set to 1 ms. The initial velocity boundary condition was determined by adding appropriate velocity to the boundary of the ultrasonic transducer to match the PIV measurement.

### Electrophysiological recording

Whole-cell patch-clamp recordings were acquired from single neurons via patch pipettes made from borosilicate glass (1.5 mm outer diameter, 0.84 mm inner diameter) pulled to resistances of 3-5 MΩ on a micropipette puller (PC-10, Narishige). Pipettes were filled with an intracellular solution containing (in mM): 135 K-Gluconate, 4 MgCl_2_, 0.2 EGTA, 3 Na_2_ATP, 0.3 Na_2_GTP, and 10 HEPES, and PH adjusted to 7.25 with KOH, 291 mOsm. The extracellular solution contained (in mM): 150 NaCl, 3 KCl, 2 CaCl_2_, 2 MgCl_2_, 20 Glucose and 10 HEPES, PH adjusted to 7.35 with NaOH, 315 mOsm. An amplifier (Axon CNS MultiClamp 700B, Molecular Devices) was used to voltage clamp neurons at a holding potential of −70 mV. Recordings were digitized (Axon CNS Digidata 1440A, Molecular Devices), and data were acquired with pClamp software (Molecular Devices). All recordings were made at room temperature (21-25 °C).

In some experiments, calcium-free external solution containing the calcium chelator EGTA (1mM) was used to test the role of calcium influx in ultrasound-evoked neurotransmitter release. To block EPSC evoked by action potentials, 1 µM tetrodotoxin (TTX) was added to the normal external solution. In other experiments, Kynurenic Acid (2 mM) was added to the normal external solution to suppress excitatory synaptic transmission. In all cases, care was taken to keep the external pH and osmolarity constant.

### Calcium imaging

To load cells with the fluorescent calcium indicator dye, fluo-4, a cover slip of neuron culture was incubated with 1.25 μM fluo-4 AM (F14201, Thermo Fisher Scientific) in Neurobasal medium A (with 2% B27 and 1% 100X Glutamax) and incubated at 37 °C in the dark for 15 min. Several (2-3 times) gentle washes with the normal external solution were used to remove unloaded fluo-4 AM before calcium imaging. The exposure out signal of the camera (Quantum 512SC) was connected to Digidata 1440A to precisely determine the timing of calcium images relative to electrophysiological signals.

### Data analysis and statistics

Data were imported into Clampfit software (Molecular Devices) and MATLAB (R2018b, MathWorks) for analysis. The frequency of synaptic events were semi-automatically analyzed using the MATLAB based SpikeTrain function from Neurasmus B.V.

## Results

### Acoustic streaming induced by low intensity pulsed ultrasound

The goal of our experiments was to characterize the effects of ultrasound on cultured mouse hippocampal neurons. For this purpose, we integrated a 25 MHz point-target-focus ultrasound transducer into an inverted microscope equipped with electrophysiology capabilities (Figure 1*A*). During the experiment, the ultrasound transducer was operated in pulse mode with a duty cycle of 5%, a pulse repetition frequency of 5 Hz, and a total duration of 20 s (Figure 1*B*). A needle hydrophone was used to measure the acoustic fields generated by such low-intensity, pulsed ultrasound (LIPUS; Fig. S1). These inputs corresponded to pressures ranging from 0.59 to 0.91 MPa and their corresponding spatial peak-pulse average intensities, I_SPPA,_ ranged from 0.58 to 1.41 W/cm^2^ (Figure 1*C*). To determine fluid shear stress exerted on cells, acoustic streaming was empirically measured by PIV (Figure 1*D*). Time-lapse images of acoustic streaming, taken from the bottom view, showed significant fluid flow around the focal region of the ultrasound transducer. The flow velocity as well as the spatial range of flow increased with ultrasound intensity, as also shown by Fig. S2. Even though the duration of individual LIPUS pulses was 10 ms and the time interval between the pulses was 200 ms, the maximum streaming velocity occurred after the ultrasound pulses had ceased, at 10.9 ms for I_SPPA_ of 0.58 W/cm^2^ and 12.5 ms for I_SPPA_ of 0.85 W/cm^2^, and the streaming flow lasted for more than 26.5 ms for I_SPPA_ of 0.58 W/cm^2^ and 200 ms for I_SPPA_ of 0.85W/cm^2^. This result indicates that the mechanical effects associated with acoustic streaming lasted much longer than the individual LIPUS pulses. Numerical modeling with COMSOL Multiphysics software revealed the predicted flow field of acoustic streaming and the distribution of acoustic pressure from a side view (Figure 1*E*). It shows that the emitted ultrasound beam focused at the cover slip on the bottom of the recording chamber, where cells are located, and was reflected there. The velocity of acoustic streaming and the acoustic pressure both were calculated to be maximal at the focal point. Further PIV measurements of the time evolution of fluid velocities at different locations and the comparison between different I_SPPA_ are shown in Fig. S2. More details about our numerical modeling results, including the acoustic pressure, acoustic body force and acoustic streaming at different I_SPPA_, are shown in Fig. S3.

**Figure 1.**
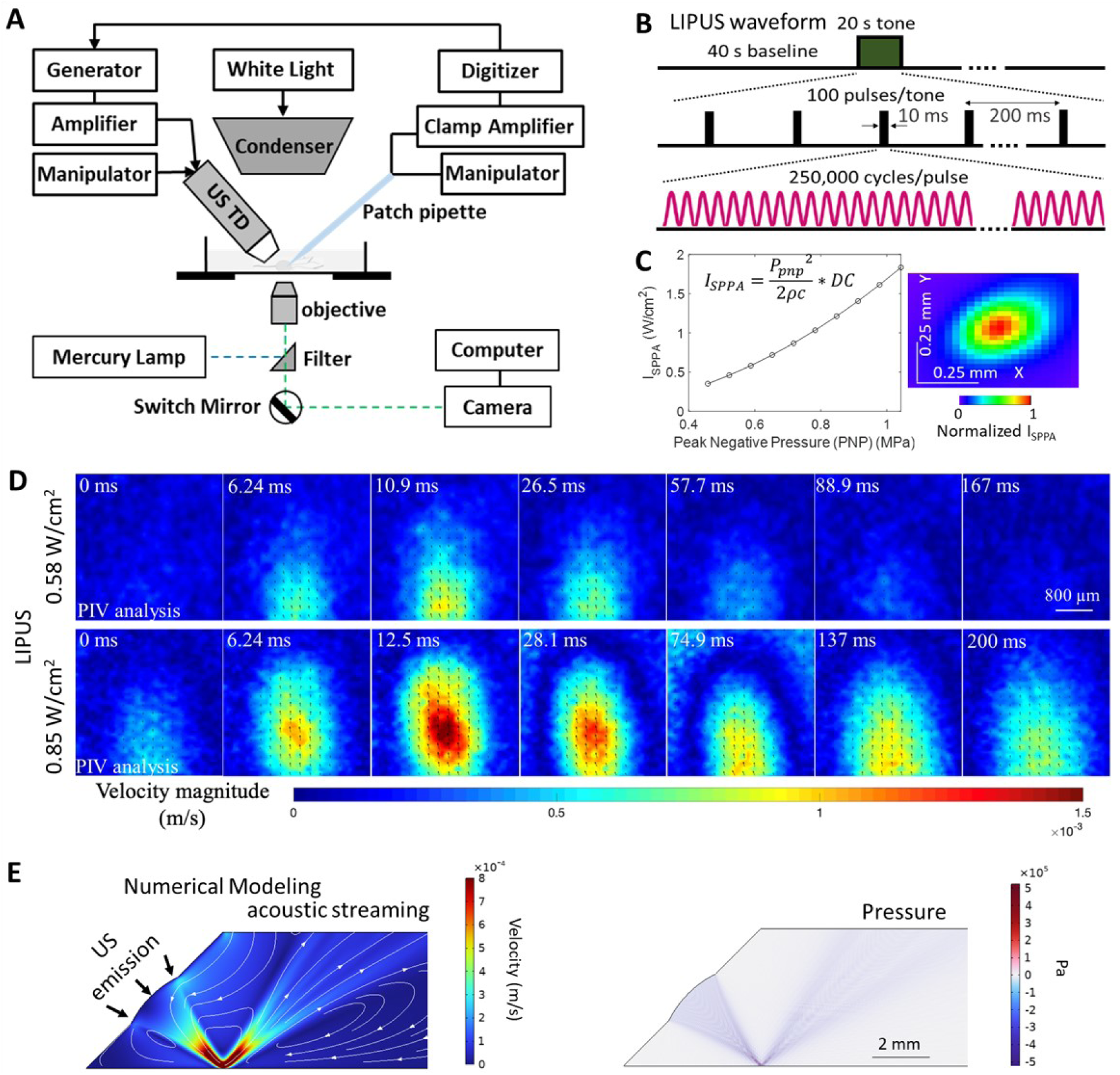
Experimental System. (A) Schematic illustration of the experimental platform for ultrasound stimulation, voltage clamp and calcium imaging of cultured neurons. A modulated sinusoidal signal was generated by a function generator, which was amplified by a 50-dB power amplifier to drive the ultrasound transducer. The position of the ultrasound transducer and the patch pipette were manipulated by three-axis micromanipulators. The timing of ultrasound stimulation and voltage clamp recordings were controlled by a digitizer. Camera was used for calcium imaging. (B) LIPUS waveform used in the experiment. 25 MHz fundamental frequency (40 ns period), pulse repetition frequency 5Hz, duty cycle (DC) 5%, total burst duration 20 s. (C) Peak negative pressure (P_pnp_) used in this study and the corresponding spatial peak pulse average intensities, I_SPPA_. Intensities were calculated using the equation shown in the inset, where ρ is the density of external solution, c is speed of sound in external solution, DC is duty cycle. (D) Time-lapsed visualization and analysis of the acoustic streaming, from bottom view, by PIV experiments done at two different ultrasound intensities (I_SPPA_ of 0.58 and 0.85 W/cm^2^). (E) Numerical simulation of the acoustic streaming and pressure distribution produced by ultrasound with COMSOL Multiphysics.

### LIPUS evokes synaptic activity in cultured neurons

We next used patch clamp recordings of ionic currents to characterize the effects of LIPUS on cultured hippocampal neurons. These recordings were made in voltage-clamp mode, with the cellular membrane potential held at −70 mV. In 20 out of 28 experiments, LIPUS application generated a barrage of apparent postsynaptic currents in neurons. Figure 2A shows an example of the high-frequency events that were evoked by LIPUS stimulation (0.46 W/cm^2^). Several lines of evidence indicate that these events are excitatory postsynaptic currents (EPSCs) produced by release of the neurotransmitter glutamate. Frist, they were abolished when LIPUS was applied in the presence of a glutamate receptor antagonist, kynurenic acid (Fig. 2B). This indicates that they are EPSCs produced by activation of ionotropic glutamate receptors. Second, they were not observed in extracellular solution containing no calcium ions (Fig. 2C). It is known that release of glutamate and other neurotransmitters requires influx of extracellular calcium [33, 34], identifying them as EPSCs resulting from glutamate release from presynaptic terminals. Third, they were abolished by treatment with tetrodotoxin (TTX, 1 μM; Fig. 2D), a well-known blocker of the voltage-dependent sodium channels that underlie neurotransmitter release evoked by presynaptic action potentials (APs). Finally, their rapid decay kinetics (< 10 ms time constant) is typical of glutamatergic EPSCs [35] (Fig. 2E). We therefore conclude that these events are a barrage of EPSCs caused by release of glutamate that is evoked by APs in presynaptic neurons that innervated the neurons that we were recording from.

**Figure 2.**
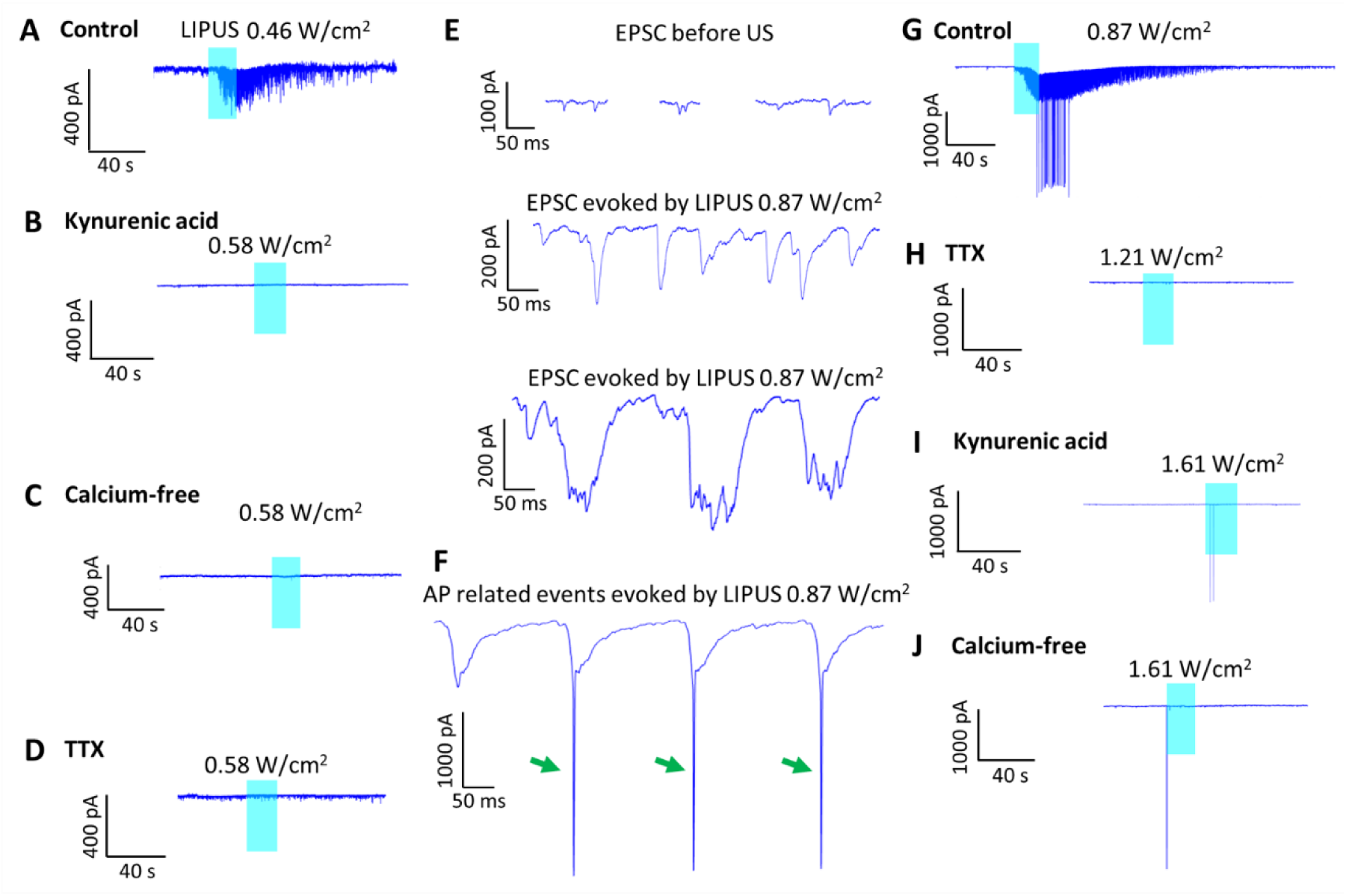
LIPUS evoked excitatory postsynaptic currents (EPSCs) in high-density neuronal cultures. Example current traces from a voltage-clamped neuron stimulated by LIPUS in normal external medium, medium with kynurenic acid (a glutamate receptor antagonist), calcium-free medium and medium with TTX at lower (A-D) and higher (G-J) ultrasound intensities. (E) Typical examples of EPSCs before ultrasound application, as well as evoked by ultrasound when there were no APs. (F) Examples of APs-related current events (arrows) evoked by ultrasound.

It is worth noting that APs are >10 times faster than EPSCs. They are also much larger in amplitude than individual EPSCs. Thus it is easy to distinguish APs from EPSCs. In some cases, the EPSCs evoked by LIPUS were large enough to evoke APs in the postsynaptic neuron, which were readily identifiable by their larger amplitudes and briefer time courses (see Fig. 2F) compared to the EPSCs (Fig. 2E). An example of such a response is shown in Fig. 2G. In this experiment, the response to LIPUS (0.46 W/cm^2^) initially consisted of a barrage of EPSCs which, in turn, generated a number of APs. Although a voltage-clamped neuron should, in principle, be prevented from firing APs, it is well-known that large depolarizations can evoke APs in distal neuronal processes whose membrane potentials are not well-controlled by the somatic voltage clamp [36, 37]. These APs were blocked by treatment with TTX (Fig. 2H). The frequency of these postsynaptic APs was reduced both by application of kynurenic acid (Fig 2I) or calcium-free conditions (Fig 2J), consistent with them arising from EPSCs. However, occasionally low-frequency AP firing could still be evoked at the onset of stronger LIPUS stimuli; we attribute these events to direct excitation of the patched neuron by LIPUS.

One striking characteristic of these responses to LIPUS was their persistence. The frequency of EPSCs gradually increased during the LIPUS stimulation and progressively decayed afterwards, with EPSCs observed for approximately a minute after the US was switched off (Figs. 3A1 and 3A2). The same was true for LIPUS-induced APs: these, too, outlasted the stimulus and, as observed for the barrage of underlying EPSCs (Figs. 3B1 and 3B2), was sustained for approximately a minute after stimulation ceased (Fig. 3B3). Thus, the mechanism responsible for the barrage of EPSCs persists for almost a minute after LIPUS.

**Figure 3.**
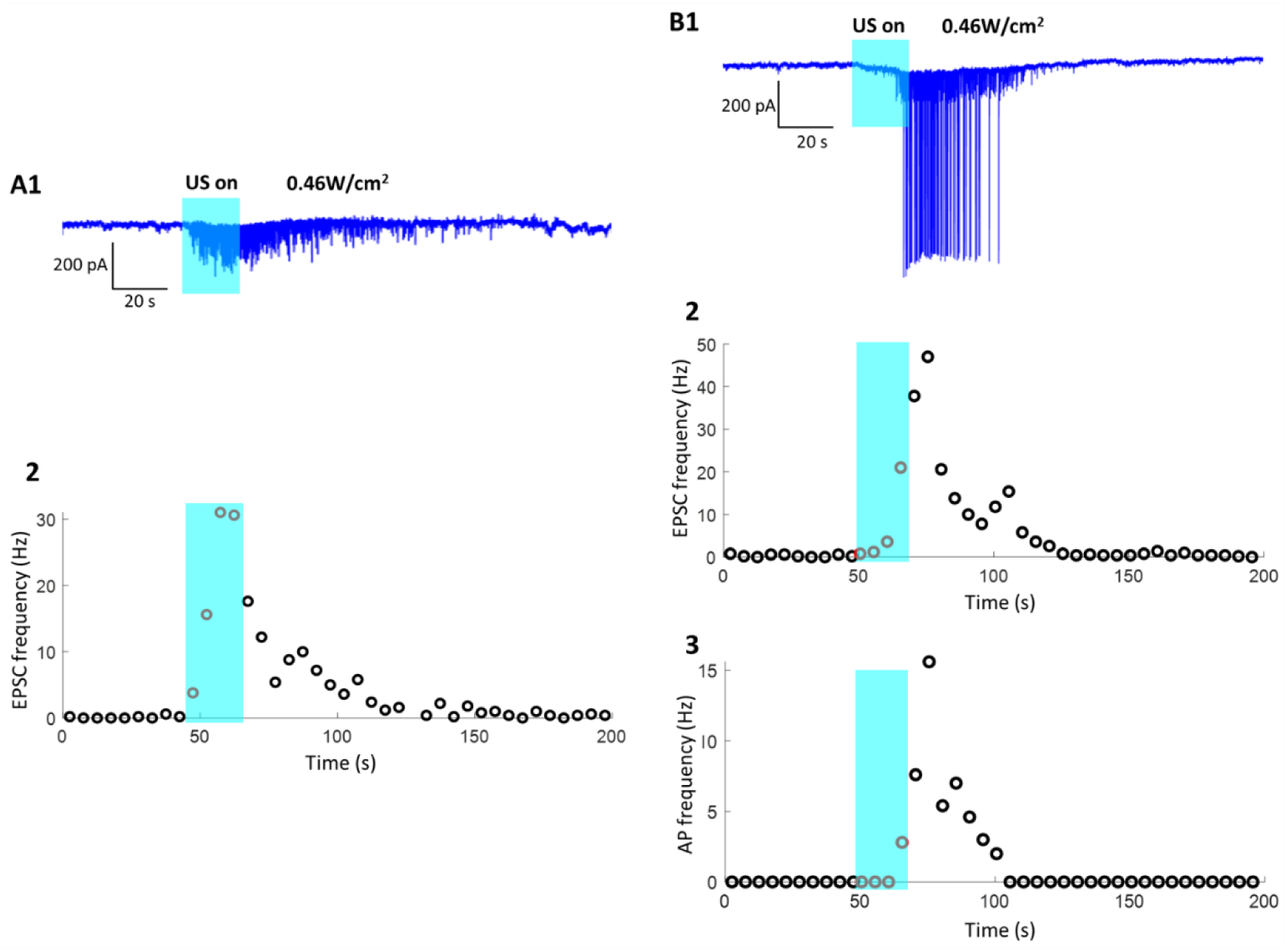
Ultrasound (blue shading) evoked sustained excitatory postsynaptic currents (EPSCs) and action potentials (APs) in high-density hippocampal neuron cultures. (A1-B1) Examples of current responses to ultrasound stimulation (0.46 W/cm^2^) in two independent experiments (both cells held at −70 mV). (A2) Time course of changes in EPSC frequency measured in the experiment shown in A1. (B2-B3) Time course of LIPUS-induced changes in EPSCs frequency (B2) and AP frequency (B3), measured in the experiment shown in B1.

We next examined the effects of different ultrasound intensities on the response of the cultured neurons to LIPUS. Figs. 4A-C show three representative examples of current measurements made during LIPUS stimulation with increasing I_SPPA_, where individual cells were voltage-clamped at a holding potential of −70 mV. In these experiments, ultrasound intensity was gradually increased until sustained EPSC or AP responses were evoked, with at least a 100 s interval between stimuli. The threshold level of LIPUS required to evoke such responses varied from experiment to experiment and ranged from 0.35 to 1.41 W/cm^2^. In general, the probability of evoking EPSC and APs increased as ultrasound intensity was increased: in all of the examples shown in Fig. 4, lower LIPUS intensities evoked smaller and briefer responses – consisting of fewer EPSCs and APs - than were observed when the stimulus intensity was increased. It is worth noting that once sustained EPSC or AP responses were evoked, equally robust responses could not be evoked again with the same or higher US intensities applied within several hundred seconds (see SI Fig. S4). Considering the results of all experiments (N = 28), the probability of LIPUS enhancing EPSC activity increased with ultrasound intensity, beyond an apparent threshold of around 0.35 W/cm^2^ (40% of cells were responsive at this intensity), and was maximal at approximately 1 W/cm^2^ (Fig. 4D). To characterize the magnitude of responses at different LIPUS intensities, we measured the total amount of charge (temporal integral of current) associated with the barrage of LIPUS-evoked synaptic activity. The total amount of charge evoked by medium (0.72-1.03 W/cm^2^) or higher intensity (1.21-1.61 W/cm^2^) LIPUS was significantly larger than that measured without US stimulation, while responses measured during lower levels of LIPUS (0.35-0.58 W/cm^2^) were smaller. This was true whether charge measurements included only EPSCs (Fig. 4E) or charge associated with both EPSCs and APs (Fig. 4F). We conclude that the magnitude of the synaptic response to LIPUS depends upon the intensity of the US stimulus. This relationship was abolished in cells treated with TTX, kynurenic acid, or calcium-free saline solution (Fig. 4G and Fig. S5), again indicating the central role of glutamate release at excitatory synapses in these responses.

**Figure 4.**
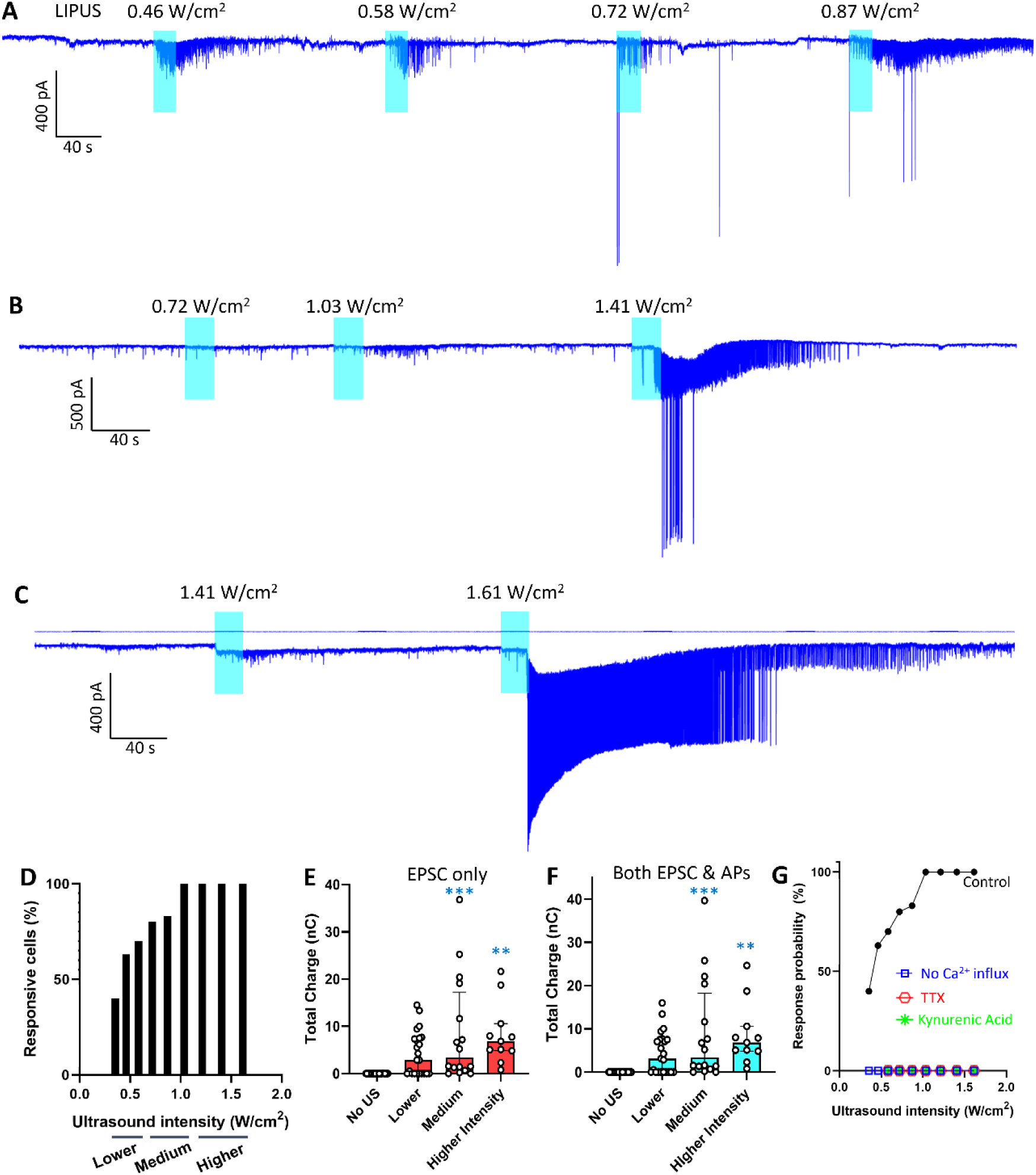
(A-C) Three different examples of current responses to LIPUS stimulation measured in high-density neuron cultures. In each case, ultrasound intensity was gradually increased until maximal sustained increases in EPSCs and/or APs occurred. Cells were held at a potential of −70 mV. (D) Effect of LIPUS intensity on the probability of evoking EPSC activity in individual neurons. N=5, 8, 10, 5, 6, 5, 5, 3, 3 for LIPUS with intensity ranging from 0.35 to 1.61 W/cm^2^, respectively. (E-F) To quantify LIPUS responses, the total charge of EPSCs alone (E) and charge of both EPSC and APs (F) was determined at different ultrasound intensities. No US group, N=24. Lower intensity LIPUS: 0.35-0.58 W/cm^2^, N=23. Medium intensity LIPUS: 0.72-1.03 W/cm^2^, N=16. Higher intensity LIPUS:1.21-1.61 W/cm^2^, N=11. The bar graph indicates the median value with the error bars denoting interquartile range; three consecutive LIPUS intensities were binned for better visualization. One-way ANOVA multiple comparisons were used for statistical comparisons; both (E) and (F) exhibited statistically significant (p<0.001) effects of LIPUS intensity. The asterisks depicted the statistical significance between specific ultrasound group and control group with no ultrasound. ** p<0.01, *** p<0.001. (G) Relationship between probability of observing LIPUS responses (enhanced EPSC frequency) and LIPUS intensity in individual neurons in normal control medium (the same control group as in Fig. 4D), calcium-free medium, medium with TTX added to block APs, and medium containing kynurenic acid to block excitatory synaptic transmission. N=2,2,9,9,9,9,7,6,5 for calcium-free medium, N=4,4,4,4,4,3,3 for medium with TTX, and N= 7,5,6,6,6,5,6 for Kynurenic Acid treated cases for the corresponding US intensities as shown in panel G.

The temperature change caused by LIPUS in our study is negligible (Fig. S6) thus not affecting the synaptic transmission. Please note that even LIPUS intensities as high as 1.61 W/cm^2^ had no effect on holding current (Fig. S7), indicating that neurons were not being damaged under our experimental conditions. We observed time-dependent reduction in AP amplitude in Fig. 4 A-C. Even though APs are all-or-none, in central neurons there is a well-known phenomenon called amplitude adaptation, that is with constant stimulus the AP gradually gets smaller over time --- it is pretty common observation in many kinds of central neurons and is well documented in neuroscience literature [38]. Besides, these APs recordings represented APs that were occurring far away from the cell body as mentioned previously.

### LIPUS response depends on neuron density

A hypothesis about the cause of the protracted responses to LIPUS comes from a detailed examination of these responses. As described above, in high-density cultures LIPUS evoked robust barrages of EPSCs and APs. Figure 5A shows another example of the response to LIPUS under these conditions: sustained EPSC activity and AP firing were observed during and following US stimulation. The inset on the right of Fig. 5A is an expanded view of part of the response, during the time that AP firing was initiated. This view reveals repetitive bursts of compound EPSCs, each of which lasts for approximately 0.2 s and attains a peak value of approximately 500 pA. Burst of EPSCs means multiple EPSC events with shorter intervals than the control condition; because of the duration of individual EPSCs, the short inter-event interval caused EPSCs to ride on top of each other, forming ‘burst of compound EPSCs’. Bursts of APs and EPSCs were well documented in neuroscience literature [39].

**Figure 5.**
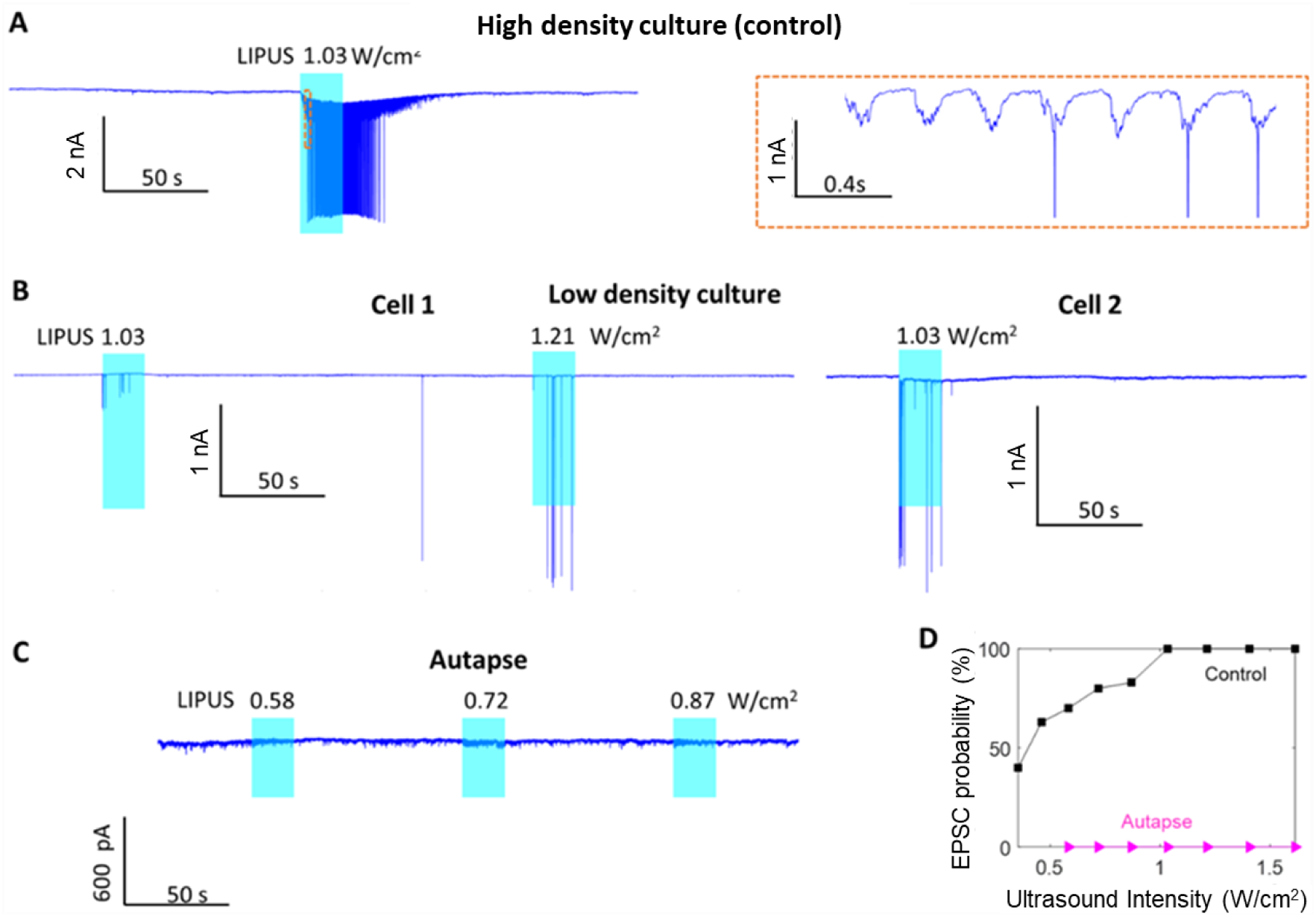
The influence of neuronal density on the ability of LIPUS to evoke EPSCs. (A) Example current response of a neuron to LIPUS in high-density culture conditions. Inset on the right shows an expanded view of part of this response. (B) Current responses to LIPUS recorded in two individual neurons in low-density culture conditions. (C) Lack of current responses to LIPUS stimulation in a microisland-cultured autaptic neuron. (D) Probability of LIPUS enhancing ESPC frequency at increasing I_SPPA_ in autaptic neurons, compared to neurons in high-density cultures (the same control group as in Fig. 4D). N=4-7 for autaptic neurons and 4-10 for the high-density control group.

As the bursts increased in size, APs were also generated. Such bursts are often observed in high-density cultures of neurons and are thought to arise from the recurrent activity of excitatory synaptic networks formed between the neurons [40, 41]. Thus, it is possible that LIPUS produces sustained EPSC and AP activity by activating such recurrent excitatory circuits.

One line of evidence supporting this possibility came from measurements made in cultures which had a lower density of neurons (∼20,000 cells/coverslip, compared to ∼80,000/coverslip for high-density cultures). In such low-density cultures, responses to LIPUS were greatly attenuated. Fig. 5B shows measurements from two neurons from different low-density cultures during LIPUS application. Even at higher ultrasound intensities (1.03 and 1.21 W/cm^2^) no bursts of EPSCs were observed. Aside from a few low-frequency EPSCs, there were a small number of APs observed, which presumably reflected LIPUS directly stimulating AP firing in these neurons. Similar results were observed in a total of 7 experiments. To further evaluate the role of neuronal density in the response to LIPUS, we also examined the effects of LIPUS in single-neuron microisland cultures, where the solitary neuron forms “autapses” with itself [32]. In 37 recordings from neurons in such conditions, no EPSC or AP activity was evoked by LIPUS (Fig. 5C). Across the entire range of LIPUS intensities, the probability of US evoking ESPCs in autaptic neurons was zero, compared to 40-100% in high-density cultures (Fig. 5D).

Taken together, our results indicate that neuronal density has an important influence on the response of cultured neurons to LIPUS. This indicates that the degree of network connectivity is a key determinant for this response and is consistent with the hypothesis that the response arises from recurrent synaptic excitation within the networks formed by the cultured neurons.

### Recurrent synaptic circuit activity underlies the response to LIPUS

The recurrent network hypothesis predicts that LIPUS should cause sustained activation of numerous neurons within the high-density cultures. We tested this prediction by using calcium imaging to monitor the activity of many neurons simultaneously. Electrical excitation of neurons is associated with transient elevation of intracellular calcium concentration, allowing fluorescent indicators to visualize calcium signals that serve as a surrogate of AP firing [42, 43]. In these experiments, we loaded neurons with the calcium indicator dye, fluo-4, and imaged many cells simultaneously by using a 10× low-magnification lens with a wide field of view. A typical example of the calcium signals produced in neurons in a high-density culture in response to LIPUS is shown in Fig. 6; the panels in Fig. 6A shows calcium signals evoked by LIPUS from fluorescence images taken at the indicated times, while Fig. 6B shows pseudo-colored surface plots that make it easier to discern the location of active neurons. Only neurons with discernable calcium signals are shown, for better visualization of the spatial propagation of calcium response. When the US was switched on at t=0, only a few cells on the top left of the field exhibited calcium responses, which are located within the ultrasound focus. This activity then gradually spreads from these cells to neurons throughout the image field by the end of the LIPUS stimulation (t = 19.8 s). Similar widespread neuronal activation in response to LIPUS was seen in a total of 26 experiments. We observed random initiation locations for the calcium responses in different independent experiments. This is consistent with the size of the ultrasound focus (around 0.4 mm in diameter; Fig. 1), indicating that all the cells in the image field were within the focal region.

**Figure 6.**
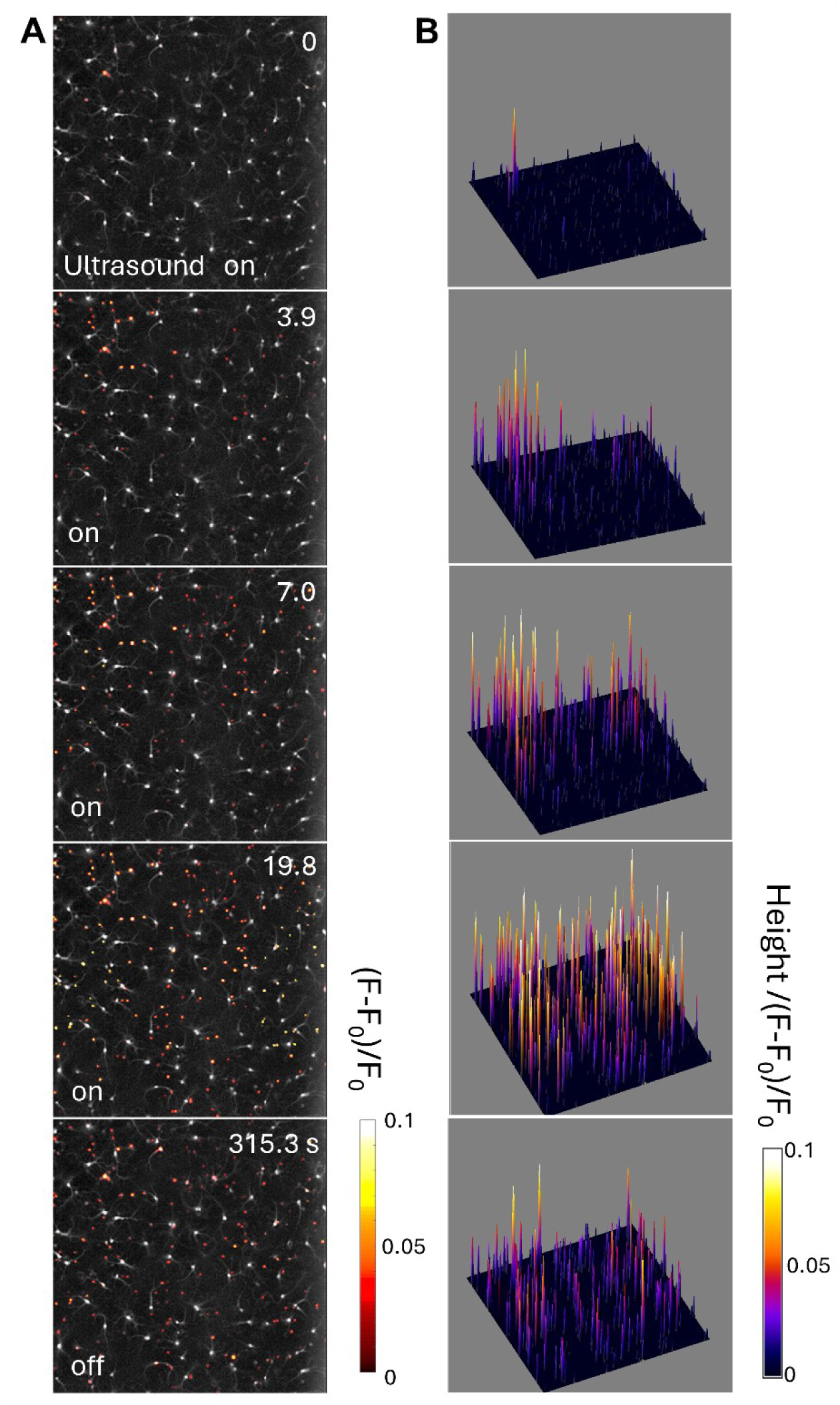
Calcium imaging of neuronal activity in a high-density culture in response to LIPUS with I_SPPA_ of 1.21 W/cm^2^. (A) Time-lapse images revealing the spatiotemporal changes in calcium signals at the indicated times after initiating LIPUS stimulation; LIPUS was applied from 0-20 s. Gray-scale image illustrates the resting fluo-4 fluorescence of all cells within the image field, while pseudocolors (scale at lower right) depict rises in calcium associated with LIPUS. (B) 3D surface plot of the calcium responses shown in (A). The height and pseudocolor (scale at lower right) indicate the magnitude of changes in calcium induced by LIPUS.

Intracellular calcium levels remained high for a minute or two after the stimulus ended; this indicates sustained activation of the neuronal network, as predicted by the hypothesis. To define the temporal relationship between neuronal activity and the EPSC barrage triggered by LIPUS, we performed simultaneous voltage clamp measurements and calcium imaging in high-density neuron cultures. Fig. 7A shows the current response of a patched neuron, while the inset shows a fluorescence image of fluo-4 loaded neurons within the image field; the location of the patch pipette is diagrammed in white and the patched neuron is indicated by a blue circle. The location of some of the neurons that exhibited calcium responses to LIPUS are labeled in red circles and the time course of their calcium responses are shown in Fig. 7B. Cells #1 to #9 are numbered in the order of their latency to produce calcium responses to LIPUS, while the calcium response of the patched cell is indicated by the blue trace in Fig. 7B. Because they are shown on the same time scale, it is possible to compare the timing of the current responses of the patched cell (Fig. 7A) with the calcium responses of cells #1 to #9 (Fig. 7B). The calcium response of cell #1 coincided with both the start of LIPUS stimulation and the barrage of EPSC activity. As more cells became active during LIPUS stimulation (Fig. 7B), the number of EPSCs increased and a few APs appeared in the patched cell (Figs. 7A,C). These APs were presumably responsible for the calcium signals observed in the patched cell (blue trace in Fig. 7B). After LIPUS was switched off, the calcium signals of the network gradually subsided; this correlated with the gradual reduction in number of EPSC events in the patched cell (Fig. 7D). The cell-averaged calcium signal of the network elevated during LIPUS stimulation, reached its peak a few seconds after LIPUS and gradually decayed afterwards (Fig. 7E). The time course of the averaged Ca^2+^ response roughly parallels that of the EPSC barrage (Fig. 7D), though it recovered somewhat more slowly than the EPSCs (see Discussion). Similar results were observed in a total of 7 experiments that showed elevated EPSC activity in the patched neurons in response to US stimulation. In 2 other experiments, we observed widespread neuronal activation that was not associated with enhanced EPSC activity in the patched cell. In these cases, it appears that the patched cell was not part of the network of activated neurons. Although the cumulative effect of LIPUS during stimulation may contribute to the intracellular Ca^2+^ response of cells, the sequential propagation of Ca^2+^ signaling and the sustained EPSC activity in the patched neuron in Fig. 6 and Fig. 7 indicate that neural network transmission is involved.

**Figure 7.**
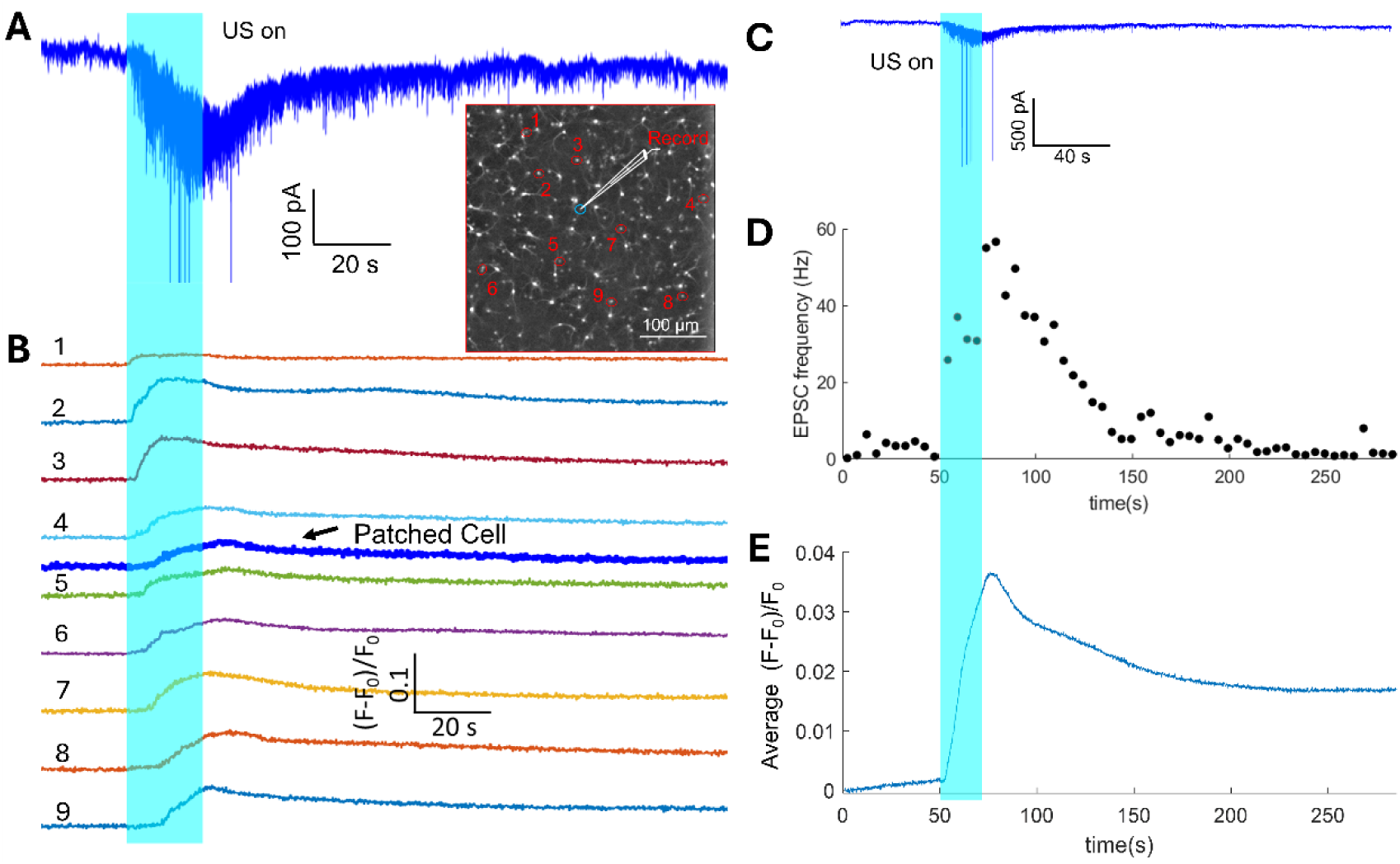
Simultaneous calcium imaging and voltage-clamp recording to compare the timing of EPSC and network responses to LIPUS with I_SPPA_ of 1.21 W/cm^2^ (applied at shaded bars). (A) Current response of an individual neuron within a high-density culture. Inset: Fluorescence image indicating location of the patch pipette (in white), with the patched cell within the blue circle. Red circles depict the location of cells exhibiting rise in calcium in response to LIPUS; cells #1 to #9 are numbered in the order which their calcium responses started. (B) Time course of calcium responses of the individual cells identified in the inset of (A), temporally aligned with the current trace shown in (A). (C-E) Comparison of time course of EPSC and calcium responses. (C) Low-gain display of the current response shown in (A), to allow visualization of APs. (D) Time course of the changes in EPSC frequency measured in the patched cell. (E) Time course of calcium response averaged over many cells in the image field.

In summary, our calcium imaging results indicate that (1) there is recurrent activation of a network of neurons in response to LIPUS; and (2) such neuronal activity is temporally coincident with the barrage of EPSCs recorded in the patched cell in response to LIPUS. Both of these findings are predicted by the hypothesis. When taken together with our observation that the barrage of EPSCs arises from APs in presynaptic neurons (Figs. 2 and 4), our results lead us to conclude that LIPUS causes the EPSC barrage via recurrent activation of an excitatory neural network within the culture.

## Discussion

Neurons in the brain are interconnected by synapses to form intricate networks, which allow the brain to perform specific functions and respond to changes in the external environment. We have found that US application increases the activity of excitatory synapses, which in turn leads to recurrent synaptic network activity that greatly outlasts the duration of US stimulation. Our results yield new insights into the actions of US on brain tissue and suggest a possible mechanism for the ability of US to modulate brain activity in vivo.

Most studies investigating the cellular mechanisms underlying US neuromodulation have employed calcium imaging [44–48], which provides good spatial resolution but provides little information about how (or whether) such calcium signaling affects neuronal function. Our electrophysiological analysis, based on whole-cell patch-clamp recording, yielded high temporal resolution and precise information about the generation of APs, as well as subthreshold events, such as EPSCs, that are not detected by calcium imaging. While several recent studies have obtained electrophysiological recordings during application of high US frequencies (30 and 43 MHz), these measurements were limited to analysis of AP dynamics in single cells and did not consider postsynaptic currents or integration of signals across neuronal networks [49–51]. Many other previous studies have focused on responses of cells expressing exogenous ion channels that are sensitive to ultrasound [49, 51, 52]. Considering the effects of US on native neurons and neuronal networks expressing only endogenous ion channels – as we have done - provides important insights that are directly relevant to US neuromodulation in vivo and offers potential insights into the use of US for clinical applications.

A recent study reported artifactual effects of US stimulation related to US-induced electrode resonance or displacement; such effects could introduce depolarizing leak currents, particularly at sub-MHz frequencies [53]. However, at the higher US frequency that we examined (25 MHz), no such displacement of the recording electrode was observed. This is consistent with other reports of successful electrophysiological recordings at high (30 and 43 MHz) US frequencies [49–51]. Displacement of the recording electrode by US could also apply mechanical stimulation to an individual neuron, which could then initiate network activity. However, this was not the case: our results shows that the patched cell was never the first one to generate calcium signals (Fig. 7B). Instead, we found that patched cells exhibited increased EPSC activity that must have arisen from LIPUS activation of other presynaptic cells. Further, in some experiments we did not use recording electrodes but still observed network calcium responses to US that were comparable to those measured in experiments that employed both calcium imaging and electrodes. Another concern is the potential for the recording electrode to introduce cavitation nuclei; this is unlikely in our case, because we used 25 MHz fundamental US frequency and the cavitation threshold is hard to reach at this high frequency with low intensity US [54]. In summary, we are confident that electrode movement was not responsible for the responses that we report here. It is worth noting that inhibition of neuronal activity by LIPUS has also been reported, e.g., for suppression of epilepsy [55, 56]. These studies usually employ ultrasound with fundamental frequency < 1MHz and PRF of 100 – 1.5 KHz, which is much larger than the PRF used in our study (5 Hz). Because activation of interneurons will inhibit synaptic networks, it is also possible that different ultrasound stimuli could preferentially activate interneurons. Finally, stimulating neurons for too long or too frequently can make them inactive, due to depolarization block [57]. Thus, whether LIPUS evokes excitatory or inhibitory effects on neuronal networks may well depend upon the ultrasound stimulus employed and this must be kept in mind in future studies. Although we used high frequency ultrasound in this study, we also observed increased calcium signaling of the neuron network for 2 min after 0.512 MHz LIPUS stimulation for 20 s (all other ultrasound parameters are the same except the fundamental frequency) by using in vivo 2-photon imaging of the mouse S1 cortex. Therefore, similar effects are likely to be evoked by LIPUS with a fundamental frequency that is suitable for in vivo study, and potentially for human therapy. One of the most important lines of evidence supporting our recurrent network model for the response to US is the observation that both the synaptic responses of individual neurons and the recurrent network activity of other neurons in the culture greatly outlast the duration of the LIPUS stimulus (Fig. 7). However, there was a quantitative mismatch between these responses: the network calcium signals (Fig. 7E) somewhat outlasted the EPSC barrage (Fig. 7D). There are several possible reasons for this mismatch. First, we would expect the EPSCs to decline more rapidly because of the non-linear (4th power) relationship between Ca^2+^ and transmitter release probability [58]. Second, it is likely that synaptic depression occurs over the prolonged time course of the response, which will deplete the synaptic vesicles and lower glutamate release even though Ca^2+^ concentration remains elevated [59]. Finally, it is possible that the cells with the most protracted Ca^2+^ signals may not be the ones that are driving the network reverberations. For these reasons, the data shown in Fig. 7 represent strong support for the recurrent network model of US stimulation. It is worth noting that a recent study reported 15 min LIPUS stimulation could facilitate neuronal activity for about 30 min in cultured hippocampal neurons, where increased frequency of both spontaneous APs and spontaneous excitatory synaptic currents (sEPSCs) as well as elevated amplitude of sEPSCs were observed [60]. Both their and our study indicated that LIPUS stimulation evokes sustained neuronal activity that persists after stimulation has stopped. It is reasonable to expect that longer stimulation would yield a more prolonged response in their study.

Recently, a study of cultured cortical neurons culture indicated that LIPUS activates specific calcium-selective mechanosensitive ion channels, resulting in a gradual build-up of calcium that is amplified by calcium- and voltage-gated channels to generate a burst-firing response [61]. Such responses are consistent with, and could even be responsible for, the recurrent excitatory neural network that we observed. Further, the recurrent network activity that we observed could be responsible for the induction of long-lasting synaptic plasticity, such as long-term potentiation or depression, by transcranial focused ultrasound [38, 62]. These long-term synaptic changes are connected to learning and memory processes that play important roles in brain function. Thus, our results provide mechanistic insights into the actions of transcranial focused ultrasound in vivo.

## Conclusions

Our results establish that LIPUS can activate neuronal network activity, EPSCs and glutamatergic synaptic transmission in high-density cultures of neurons. Such activation lasts for tens to hundreds of seconds and is enhanced by higher levels of US. Extracellular calcium influx, APs firing, synaptic transmission, neuron network connectivity and the recurrent excitatory network activity are all necessary for LIPUS-activated EPSCs. Our results provide insights into the mechanism of LIPUS-induced neuromodulation and reveal strategies for utilizing LIPUS to regulate synaptic transmission and neurotransmitter release in vivo, with the potential for intervention in neurological disorders such as Parkinson’s disease and Alzheimer’s disease.

## Abbreviations

LIPUS: low-intensity pulsed ultrasound
EPSC: excitatory postsynaptic current
AP: action potential
TTX: tetrodotoxin
PD: Parkinson’s disease
AD: Alzheimer’s disease
PIV: particle image velocimetry
CFD: computational fluid dynamic.

## Acknowledgments

The authors thank Prof. Yufeng Zhou from Nanyang Technological University and Chongqing Medical University for providing equipment and guidance for the X-Y-Z acoustic output measurement of the US transducer, and Prof. Long Meng for the generous help of qualitative 3D mapping of US beam profile. The authors thank Melissa Yeow Wei Bao, Minchuan Zhang, Chon U Chan and Vicky Liao for technical support, and Kai Voges for the generous gift of MATLAB based SpikeTrain function for EPSC analysis. This work was supported by the National Natural Science Foundation of China, Science Foundation for Youths, 12204322, and the LKCMedicine Dean’s Postdoctoral Fellowship Grant from Nanyang Technological University, Singapore.

## Supporting Information

### Output characterization of the ultrasound transducer

The transducer output was characterized by a needle hydrophone (ONDA HNA-0400) measurements with 0.05-mm increments in lateral directions, and 0.25 mm increments in axial direction in a large tank filled with water (Fig. S1 A-D). After finding the focal point, the output peak negative pressure was measured there at different input voltage (Fig. S1 E-F). The −6 dB focal region at the lateral direction is around 0.36 × 0.39 mm while the −6 dB axial focal width is around 5mm. We further measured the acoustic beam profile generated by the transducer with a 3D motorized scanner with another needle hydrophone (ONDA HNR-0500). The normalized acoustic intensity and focal point at lateral and axial direction is shown in Fig. S1 H, which is consistent with the result from X-Y-Z scan.

### Particle image velocimetry (PIV) measurement of acoustic streaming at different ultrasound intensities

Based on the PIV flow mapping shown in Fig. 1E, we further quantified the flow velocity at specific locations at different LIPUS intensities. The measured peak flow velocity at point A, B, C along the axis of the ultrasound emission direction (Fig. S2 A) at LIPUS intensity of 0.58 and 0.85 W/cm^2^ is shown in Fig. S2 B-D, which induced by 0.85 W/cm^2^ LIPUS are significantly larger than those at lower ultrasound intensity of 0.58 W/cm^2^. Fig. S2 E-F shows the time evolution of the transient flow velocity at point A, B, C evoked by LIPUS with I_SPPA_ of 0.58 and 0.85 W/cm^2^, respectively. Given that the individual pulse duration of LIPUS is 10 ms and the time interval between the pulses is 200 ms, the velocity-time curves indicate the mechanical effects from the acoustic streaming last much longer than the individual pulse duration of LIPUS.

Similarly, the measured peak flow velocity at point D, E, F along the line perpendicular to the axis of the ultrasound emission direction (Fig. S2 G) at LIPUS intensity of 0.58 and 0.85 W/cm^2^ is shown in Fig. S2 H-J. Again, the velocities at these points induced by 0.85 W/cm^2^ LIPUS are significantly larger than those at lower ultrasound intensity of 0.58 W/cm^2^. Fig. S2 K-L shows the time evolution of the transient flow velocity at these points, which also indicates the mechanical effects from the acoustic streaming last much longer than the individual pulse duration of LIPUS. Furthermore, point E located at the center of the line demonstrates the largest velocity among these points at both ultrasound intensities.

### COMSOL Multiphysics modeling of LIPUS induced physical effects at different ultrasound intensities

Using COMSOL Multiphysics modeling as described in the main text, we get the results for the side view of LIPUS induced pressure distribution, acoustic body force and acoustic streaming at two different ultrasound intensities. The physical effects, including pressure distribution, acoustic body force and acoustic streaming, are all stronger at I_SPPA_ of 0.85 W/cm^2^ (Fig. S3 A) compared to I_SPPA_ of 0.58 W/cm^2^ (Fig. S3 B).

### Stable electrophysiological recordings at higher ultrasound frequencies

We utilized LIPUS with a fundamental frequency of 25 MHz to stimulate cultured neurons during voltage clamp recording, where stable current traces has been achieved. It is worth noting that under the same acoustic pressure, it is easier to obtain stable recording from whole-cell patch clamp at 25 MHz ultrasound compared to that at 2.25 MHz. With the latter, we could only get stable recording at very low acoustic pressure and the current trace is propone to the mechanical vibration caused by ultrasound resulting in the lost of the sealing of the pipette and the drop off of the current trace. Our result is consistent with previous studies showing stable patch-clamp recordings at higher ultrasound frequency (30 and 43 MHz).

**Figure S1.**
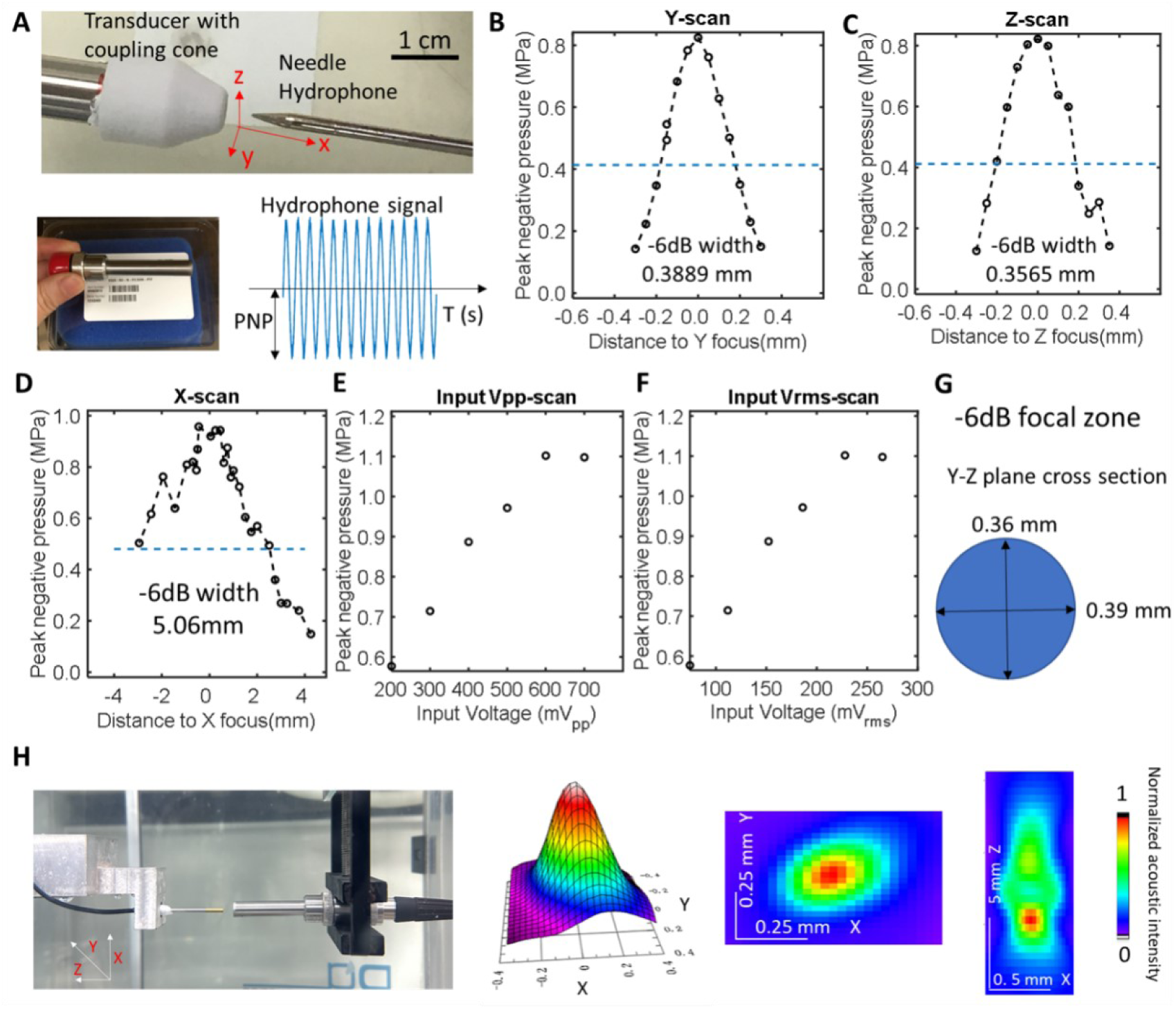
Measurement of the acoustic output of the 25MHz ultrasound transducer. (A) A sharp needle hydrophone (ONDA HNA-0400) was used to measure the pressure output of the transducer with 0.05-mm increments in lateral directions (Y-Z axis), and 0.25 mm increments in axial direction (X axis) in a large tank filled with water. (B) Obtained peak negative pressure from the Y-scan. The dashed line depicts −6 dB of the maximum pressure measured in Y-scan. (C) Obtained peak negative pressure from the Z-scan. The dashed line depicts −6 dB of the maximum pressure measured in Z-scan. (D) Obtained peak negative pressure from the X-scan (axial direction). The dashed line depicts −6 dB of the maximum pressure measured in Z-scan. After finding the focal point from the above X-Y-Z scan, the input peak-peak voltage V_pp_ (E) and the input root-mean-square voltage V_rms_ (F) of the function generator was varied to scan the output peak negative pressure. (G) Schematic showing the −6 dB focal region at the lateral direction, which is around 0.36 × 0.39 mm. (H) We further measured the acoustic field generated by the transducer with a 3D motorized scanner with another needle hydrophone (ONDA HNR-0500) in water tank. The color maps shows the normalized acoustic intensity and distribution in X-Y (lateral) and X-Z (axial) plane, from which we can infer the the focal region.

**Figure S2.**
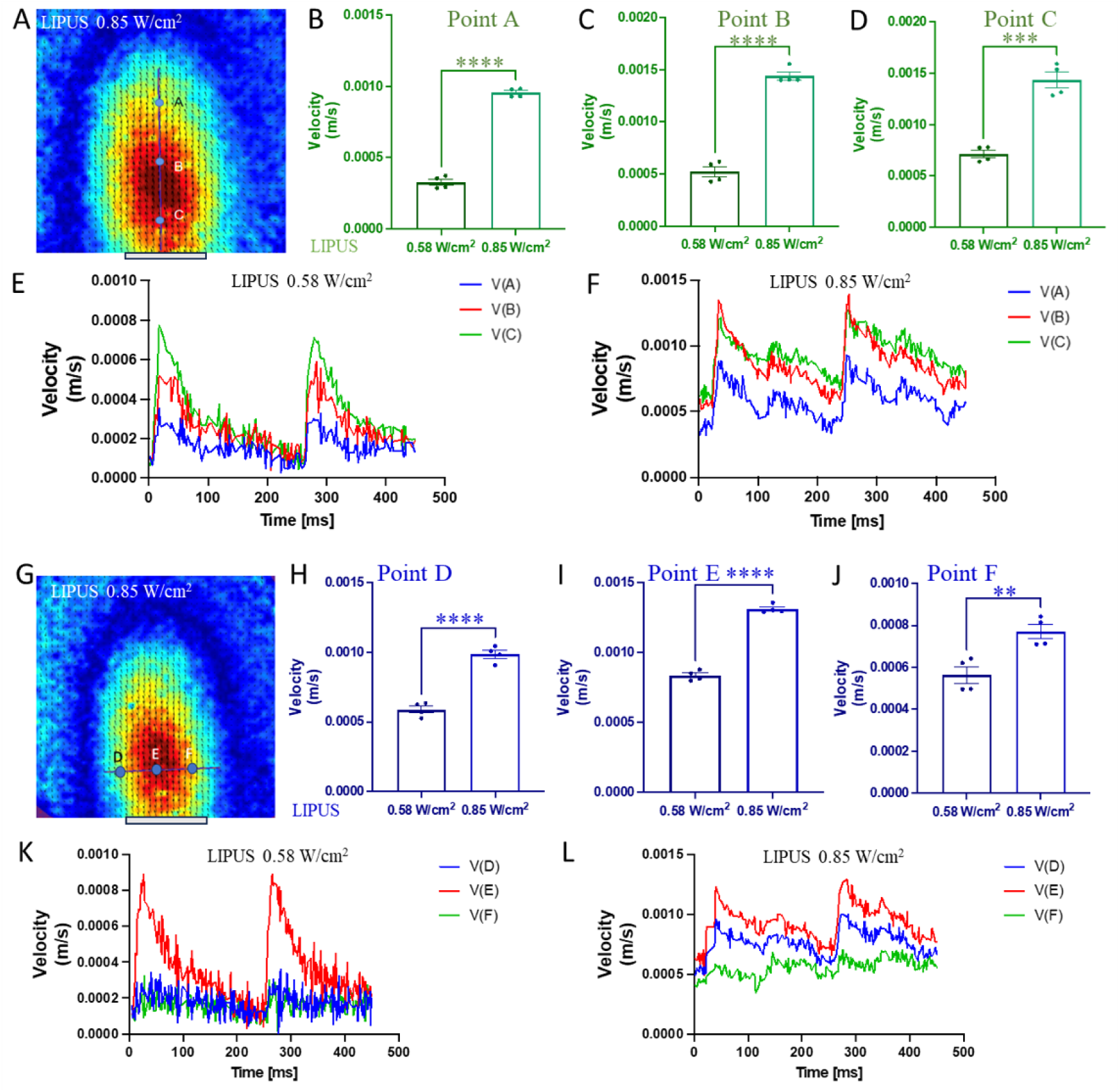
PIV measurement of the acoustic streaming induced by LIPUS in this study. (A) PIV results of the fluid flow caused by 0.85 W/cm^2^ LIPUS. A line along the axis of the ultrasound emission direction covering the flow pattern of the maximum acoustic streaming from the bottom view is dawn, where point A,B,C is located at its 0.2, 0.5, 0.8 node, respectively. The gray bar below the image depicts the emission surface of the ultrasound transducer. (B-D) The measured peak flow velocity at point A, B, C at LIPUS intensity of 0.58 and 0.85 W/cm^2^. (E-F) The time evolution of the transient flow velocity at point A, B, C evoked by LIPUS with I_SPPA_ of 0.58 and 0.85 W/cm^2^. (G) A line perpendicular to the axis of the ultrasound emission direction covering the flow pattern of the maximum acoustic streaming from the bottom view is dawn, where point D, E, F is located at its 0.2, 0.5, 0.8 node, respectively. The gray bar below the image depicts the emission surface of the ultrasound transducer. (H-J) The measured peak flow velocity at point D, E, F at LIPUS intensity of 0.58 and 0.85 W/cm^2^. (K-L) The time evolution of the transient flow velocity at point D, E, F evoked by LIPUS with I_SPPA_ of 0.58 and 0.85 W/cm^2^.

**Figure S3.**
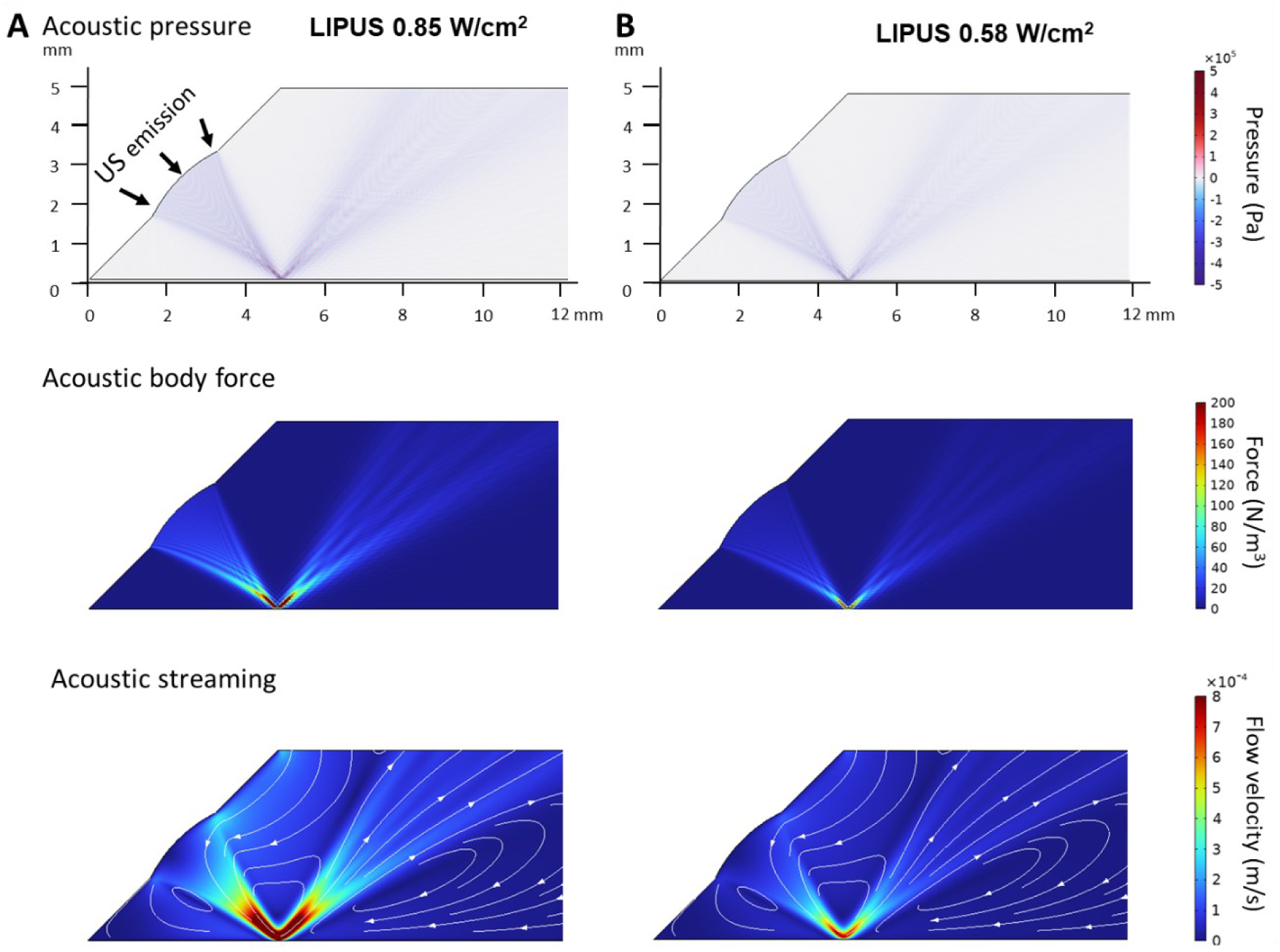
Side view of LIPUS induced pressure distribution, acoustic body force and acoustic streaming at two different ultrasound intensities from COMSOL Multiphysics modeling. (A) LIPUS at I_SPPA_ of 0.85 W/cm^2^. (B) LIPUS at I_SPPA_ of 0.58 W/cm^2^.

**Figure S4.**
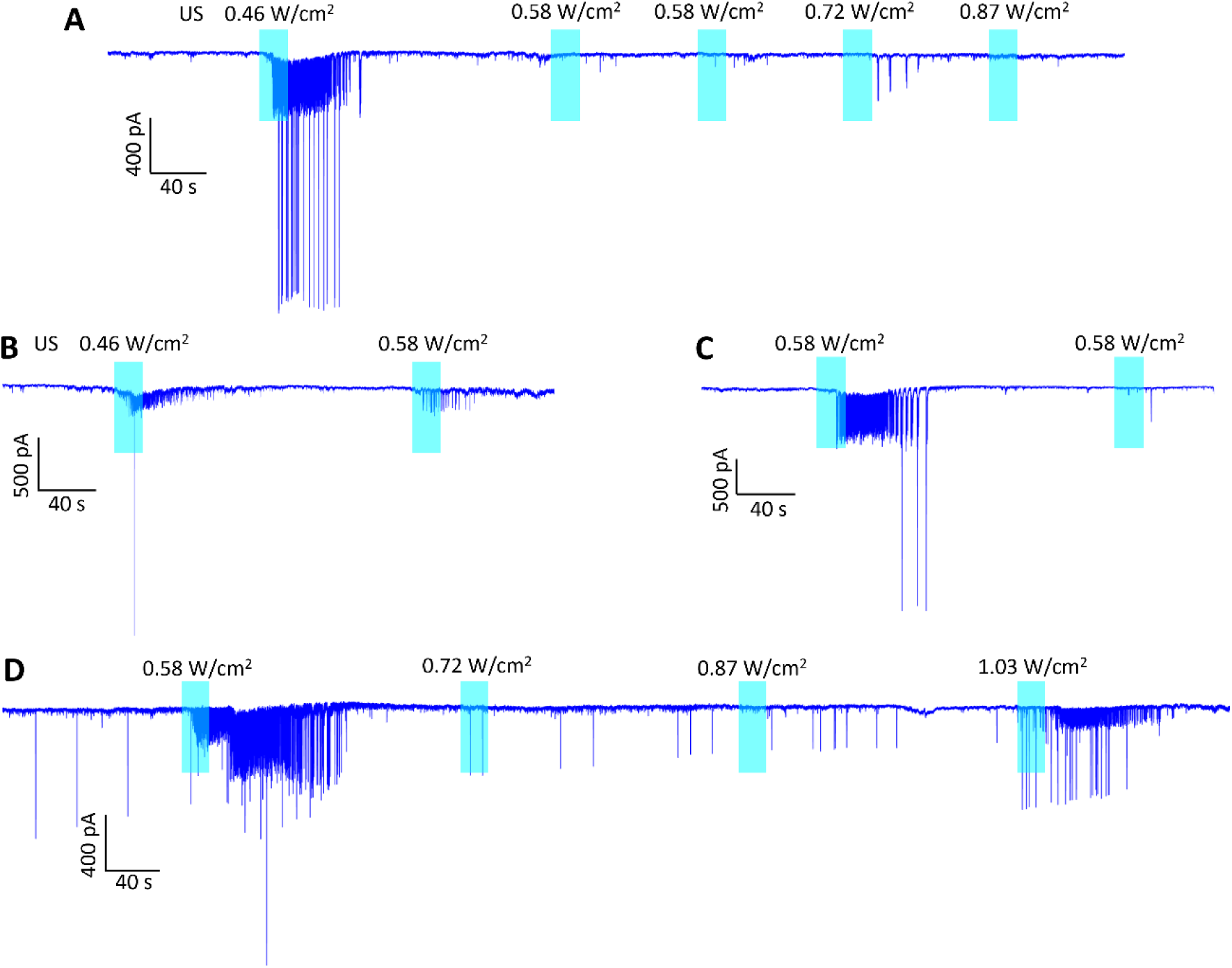
Exemplary voltage-clamp traces showing that once sustained EPSC or AP responses were evoked, this kind of response could not be evoked again with the same or higher US intensities within several 100 s.

**Figure S5.**
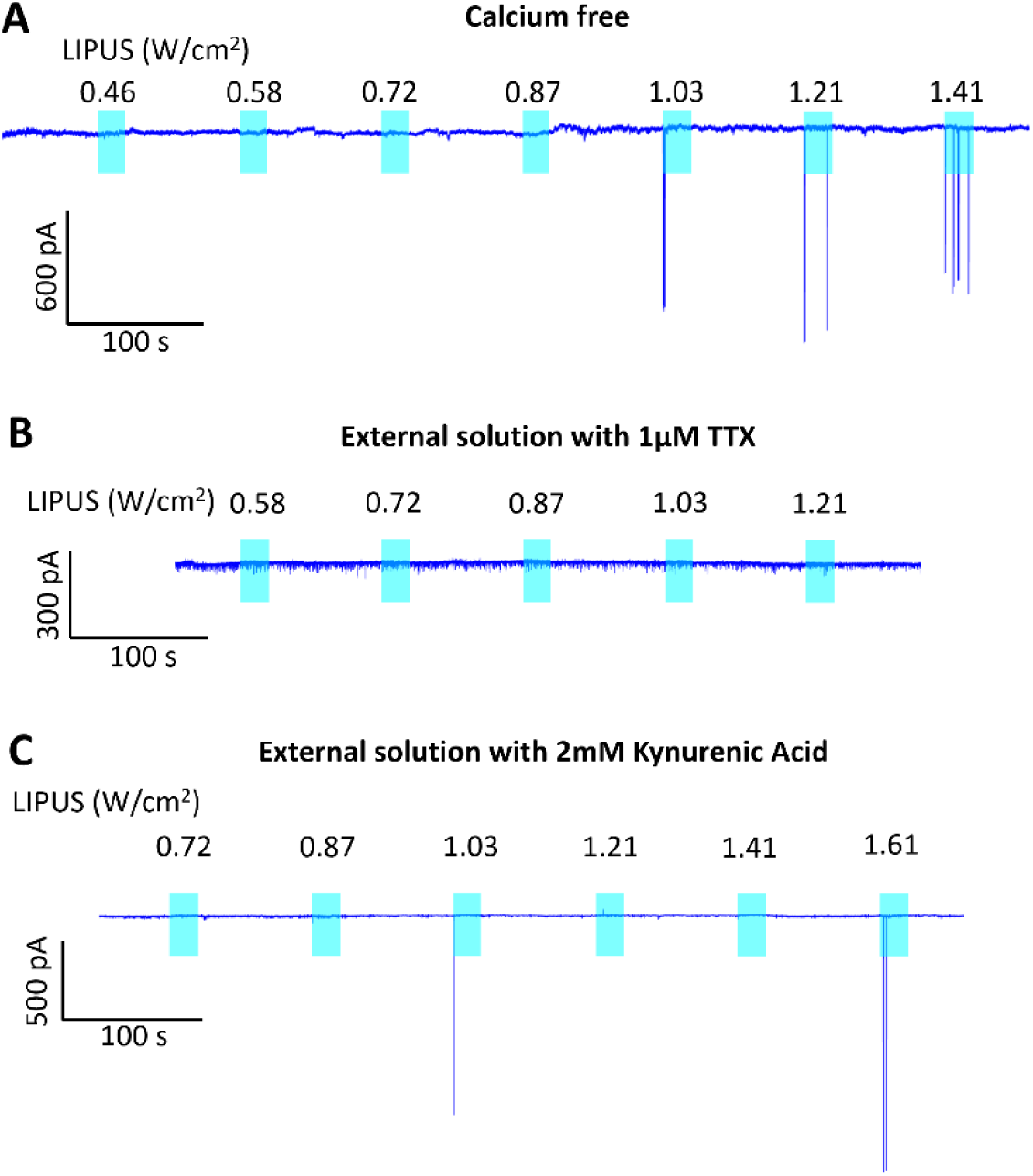
Mechanistic study in high density neuron culture. (A) Exemplary current trace for a neuron treated with calcium free external solution (ES) with 1mM EGTA and stimulated by ultrasound with increasing intensities. (B) Typical current trace for a neuron treated with 1 μM TTX in ES to block APs and stimulated by ultrasound with increasing intensities. (E) Exemplary current trace for a neuron treated with 2 mM Kynurenic Acid in ES to block synaptic transmission and stimulated by ultrasound with climbing intensities.

**Figure S6.**
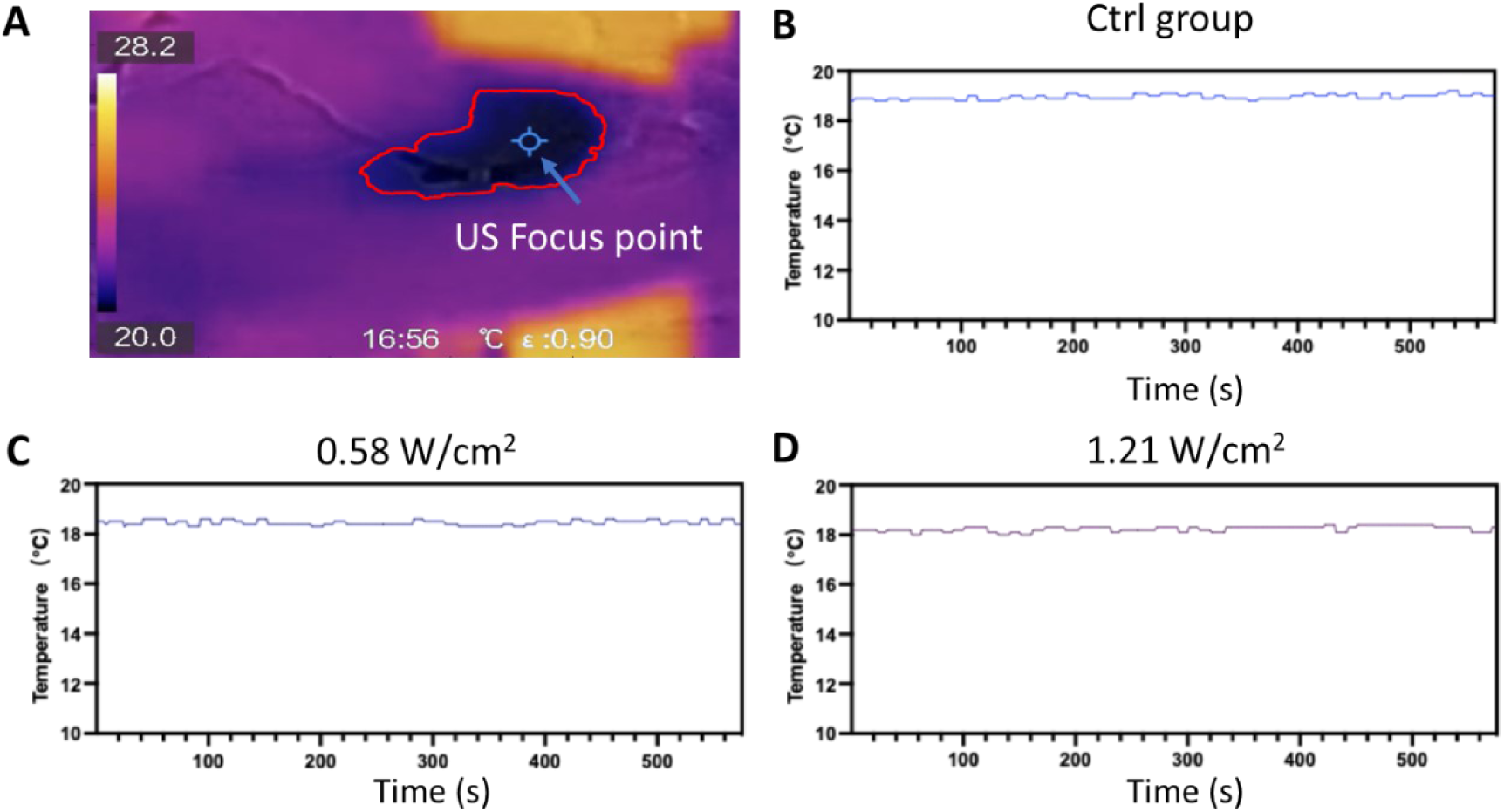
Temperature measurement. (A) *In vitro* infrared thermography (HIKMICRO, HM-TPK20-3AQF/W). The region in red outline indicates the liquid reservoir while the blue label depicts the focal region of LIPUS. (B-D) Measured temperature change over 23 seconds recording with thermography for control group without LIPUS (Ctrl) and LIPUS groups at intensities of 0.58 and 1.21 W/cm^2^.

**Figure S7.**
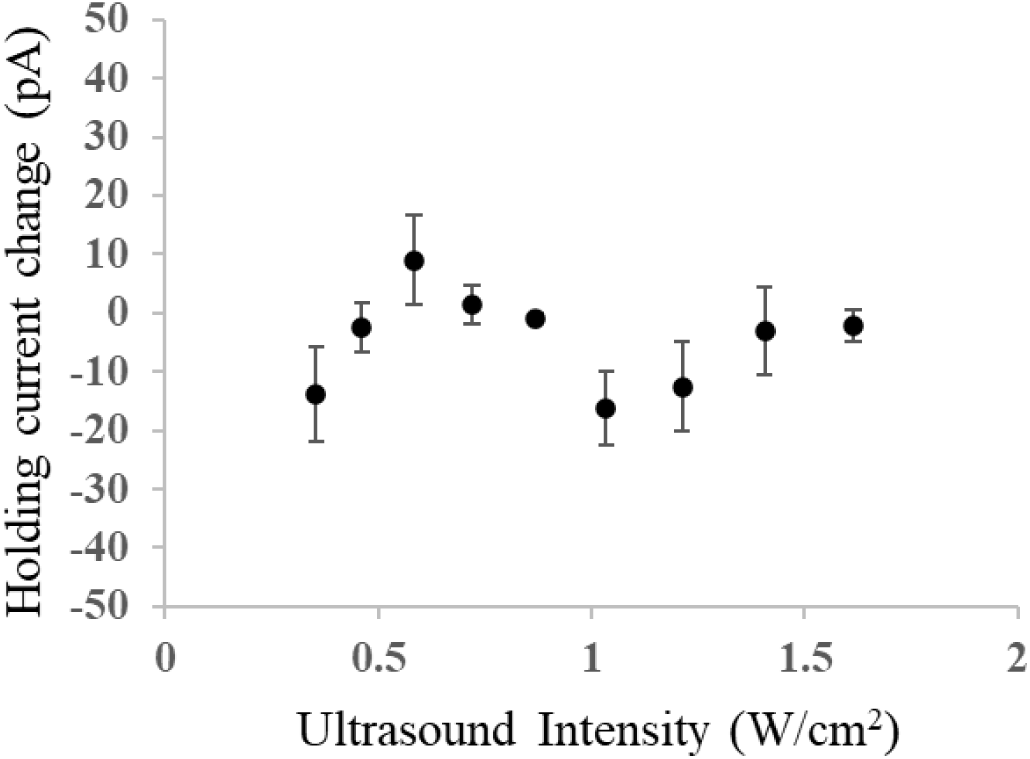
Measurement of the holding current change at different ultrasound intensities in normal medium. The change is calculated as: the holding current after LIPUS induced EPSCs have eased – the holding current before LIPUS stimulation. A positive value indicates the current trace becomes less leaky while the negative values imply slightly more leaky holding current. Data are presented as mean ± SEM. Overall, the change of the holding current is small indicating LIPUS stimulation not causing damage to the neurons.

