## Supplemental files for "Low intensity pulsed ultrasound activates excitatory synaptic networks in cultured hippocampal neurons"

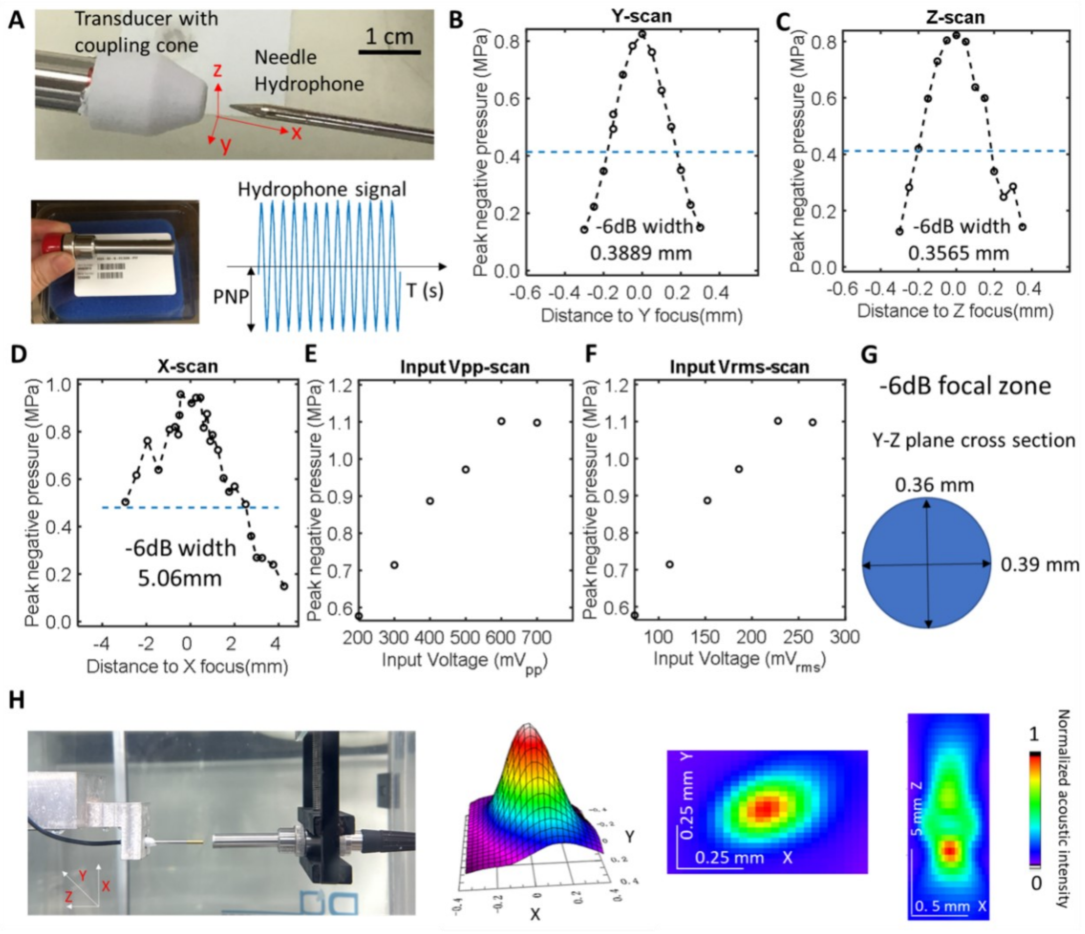


**Figure S1.** Measurement of the acoustic output of the 25MHz ultrasound transducer. (A) A sharp needle hydrophone (ONDA HNA-0400) was used to measure the pressure output of the transducer with 0.05-mm increments in lateral directions (Y-Z axis), and 0.25 mm increments in axial direction (X axis) in a large tank filled with water. (B) Obtained peak negative pressure from the Y-scan. The dashed line depicts -6 dB of the maximum pressure measured in Y-scan. (C) Obtained peak negative pressure from the Z-scan. The dashed line depicts -6 dB of the maximum pressure measured in Z-scan. (D) Obtained peak negative pressure from the X-scan (axial direction). The dashed line depicts -6 dB of the maximum pressure measured in Z-scan. After finding the focal point from the above X-Y-Z scan, the input peak-peak voltage $V_{pp}$ (E) and the input root-mean-square voltage $V_{rms}$ (F) of the function generator was varied to scan the output peak negative pressure. (G) Schematic showing the -6 dB focal region at the lateral direction, which is around 0.36 × 0.39 mm. (H) We further measured the acoustic field generated by the transducer with a 3D motorized scanner with another needle hydrophone (ONDA HNR-0500) in water tank. The color maps shows the normalized acoustic intensity and distribution in X-Y (lateral) and X-Z (axial) plane, from which we can infer the the focal region.

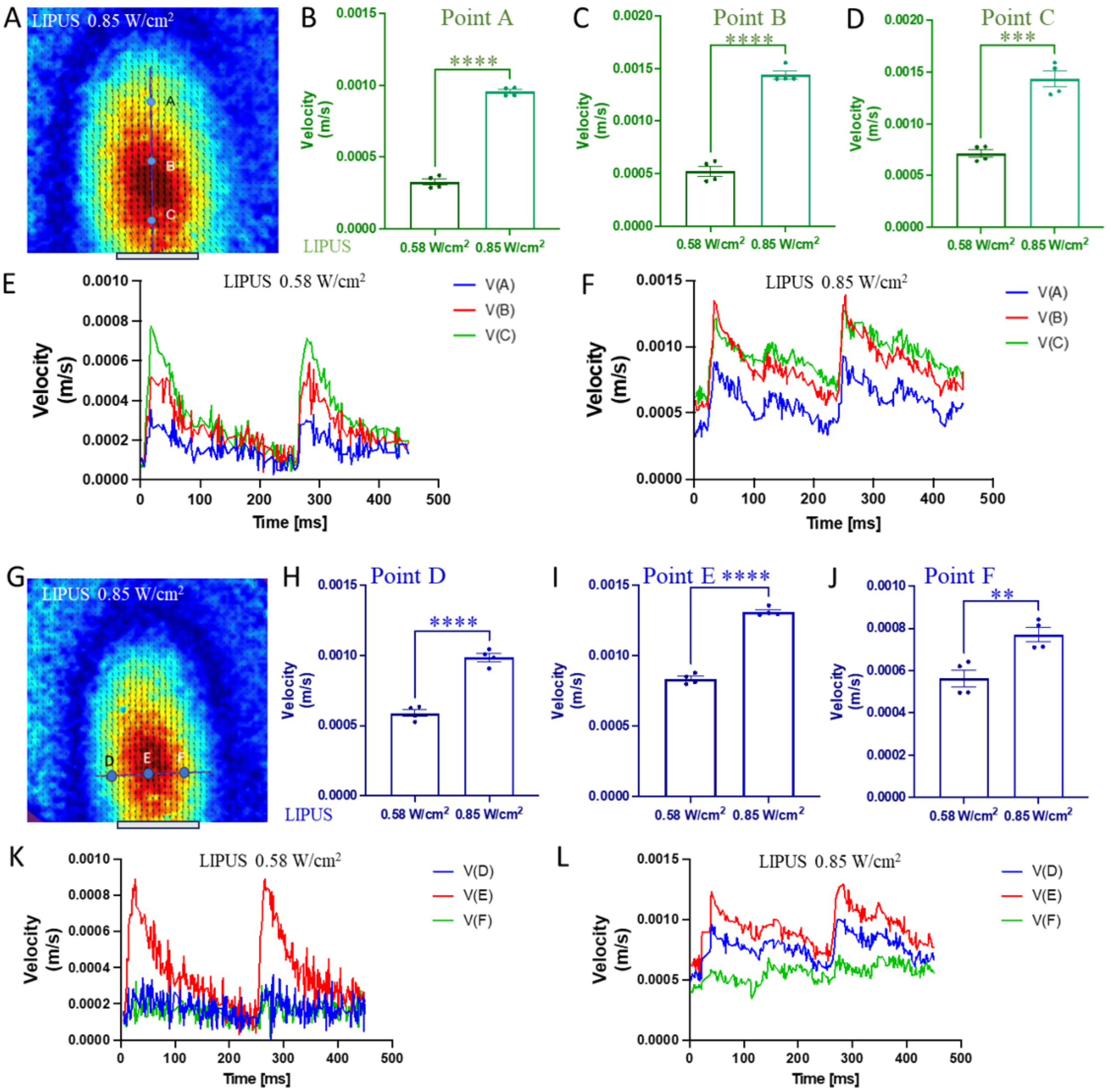


**Figure S2.** PIV measurement of the acoustic streaming induced by LIPUS in this study. (A) PIV results of the fluid flow caused by 0.85 W/cm$^2$ LIPUS. A line along the axis of the ultrasound emission direction covering the flow pattern of the maximum acoustic streaming from the bottom view is dawn, where point A,B,C is located at its 0.2, 0.5, 0.8 node, respectively. The gray bar below the image depicts the emission surface of the ultrasound transducer. (B-D) The measured peak flow velocity at point A, B, C at LIPUS intensity of 0.58 and 0.85 W/cm$^2$. (E-F) The time evolution of the transient flow velocity at point A, B, C evoked by LIPUS with $I_{SPPA}$ of 0.58 and 0.85 W/cm$^2$. (G) A line perpendicular to the axis of the ultrasound emission direction covering the flow pattern of the maximum acoustic streaming from the bottom view is dawn, where point D, E, F is located at its 0.2, 0.5, 0.8 node, respectively. The gray bar below the image depicts the emission surface of the ultrasound transducer. (H-J) The measured peak flow velocity at point D, E, F at LIPUS intensity of 0.58 and 0.85 W/cm$^2$. (K-L) The time evolution of the transient flow velocity at point D, E, F evoked by LIPUS with $I_{SPPA}$ of 0.58 and 0.85 W/cm$^2$.

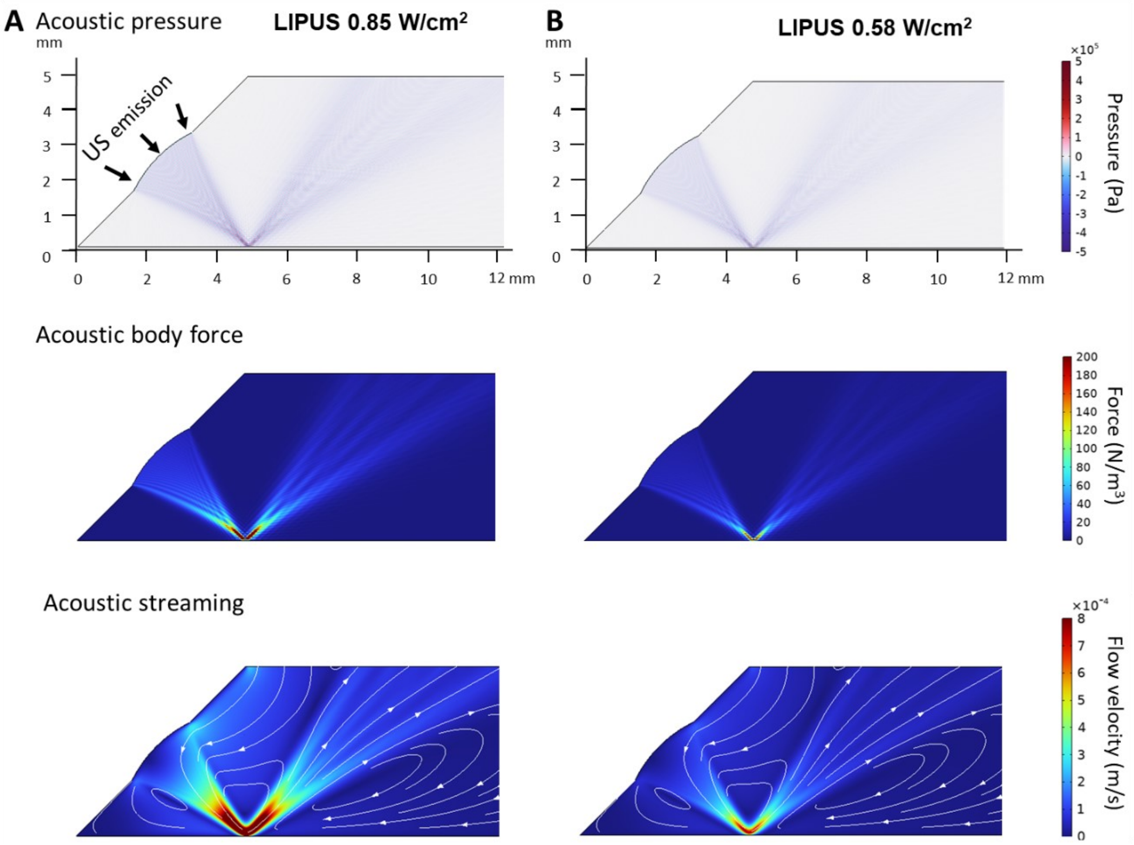


**Figure S3.** Side view of LIPUS induced pressure distribution, acoustic body force and acoustic streaming at two different ultrasound intensities from COMSOL Multiphysics modeling. (A) LIPUS at $I_{SPPA}$ of 0.85 W/cm$^2$. (B) LIPUS at $I_{SPPA}$ of 0.58 W/cm$^2$.

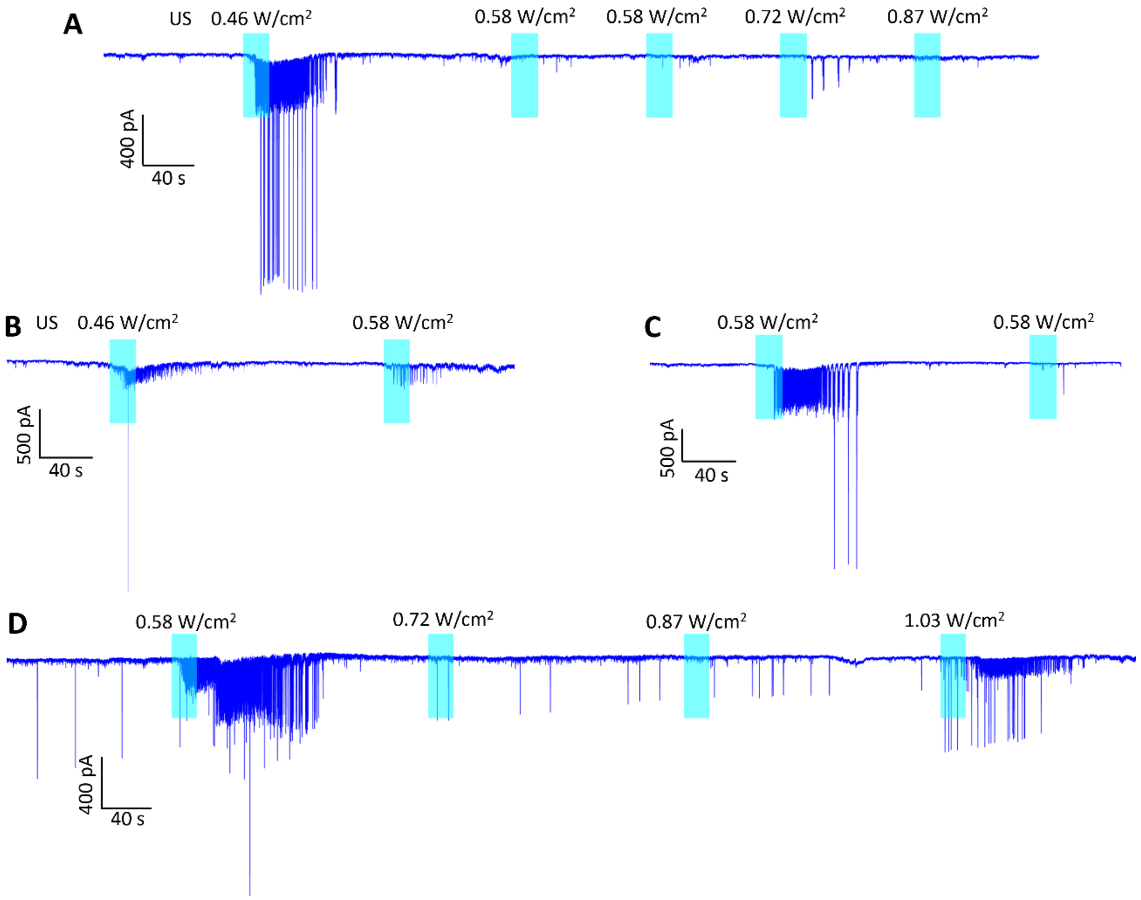


**Figure S4.** Exemplary voltage-clamp traces showing that once sustained EPSC or AP responses were evoked, this kind of response could not be evoked again with the same or higher US intensities within several 100 s.

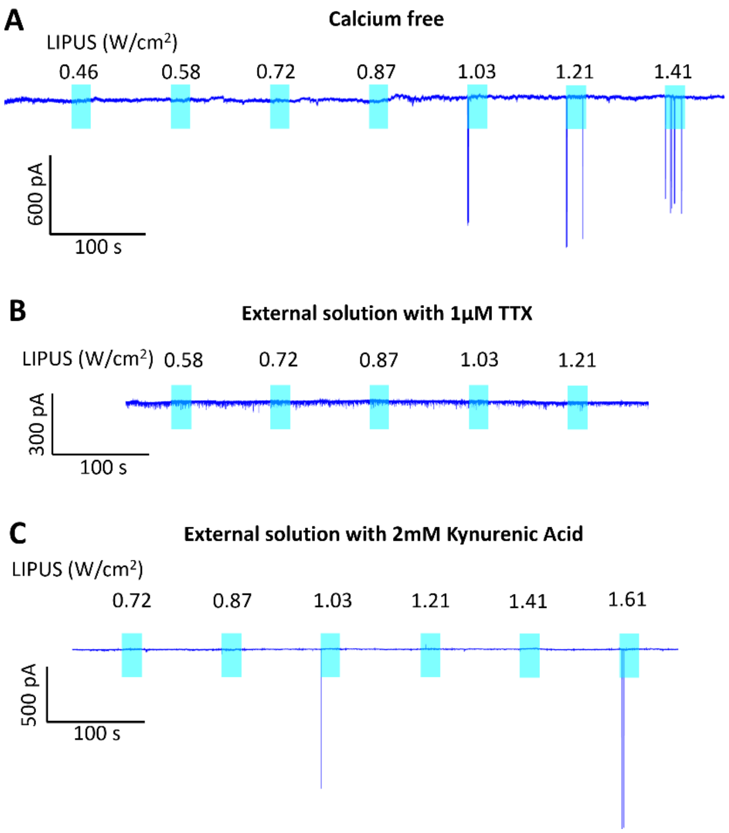


**Figure S5.** Mechanistic study in high density neuron culture. (A) Exemplary current trace for a neuron treated with calcium free external solution (ES) with 1mM EGTA and stimulated by ultrasound with increasing intensities. (B) Typical current trace for a neuron treated with 1 μM TTX in ES to block APs and stimulated by ultrasound with increasing intensities. (E) Exemplary current trace for a neuron treated with 2 mM Kynurenic Acid in ES to block synaptic transmission and stimulated by ultrasound with climbing intensities.

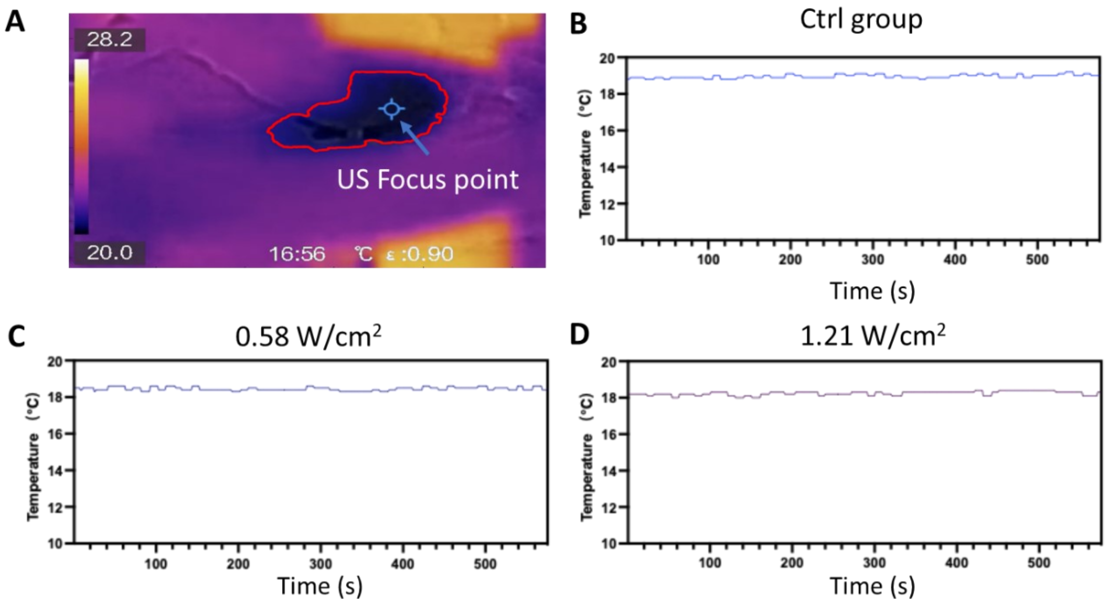


**Figure S6.** Temperature measurement. (A) *In vitro* infrared thermography (HIKMICRO, HM-TPK20-3AQF/W). The region in red outline indicates the liquid reservoir while the blue label depicts the focal region of LIPUS. (B-D) Measured temperature change over 23 seconds recording with thermography for control group without LIPUS (Ctrl) and LIPUS groups at intensities of 0.58 and 1.21 W/cm$^2$.

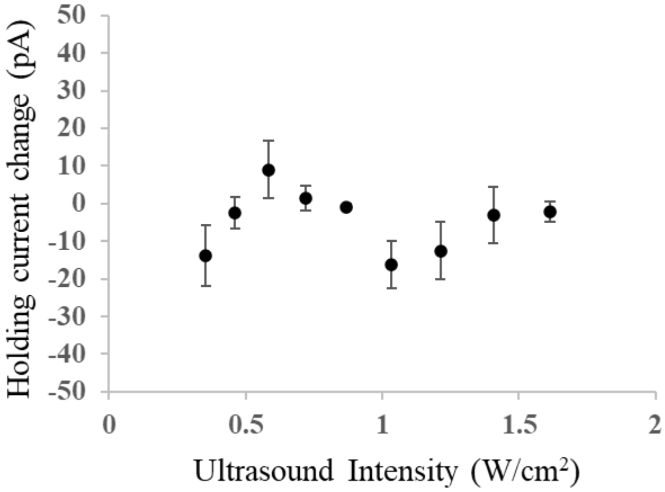


**Figure S7.** Measurement of the holding current change at different ultrasound intensities in normal medium. The change is calculated as: the holding current after LIPUS induced EPSCs have eased – the holding current before LIPUS stimulation. A positive value indicates the current trace becomes less leaky while the negative values imply slightly more leaky holding current. Data are presented as mean ± SEM. Overall, the change of the holding current is small indicating LIPUS stimulation not causing damage to the neurons.
